# RNA toehold switch-based reporter assay to assess bacterial uptake of antisense oligomers

**DOI:** 10.1101/2024.10.25.620056

**Authors:** Paramita Sarkar, Linda Popella, Sandra Pérez-Jiménez, Jörg Vogel

**Affiliations:** Institute for Molecular Infection Biology (IMIB), Faculty of Medicine, University of Würzburg, 97080 Würzburg, Germany; Cluster for Nucleic Acid Therapeutics Munich (CNATM), Munich, Germany; Helmholtz Institute for RNA-based Infection Research (HIRI), Helmholtz Centre for Infection Research (HZI), 97080 Würzburg, Germany

**Keywords:** antisense oligomers, reporter assay, peptide nucleic acids, cell-penetrating peptides, alternative antibiotics

## Abstract

Antisense oligomers (ASOs) hold promise as antibiotics for selective targeting of bacterial pathogens and as tools for the modulation of gene function in genetically intractable microbes. However, their efficient delivery across the complex bacterial envelope remains a major challenge. There are few methods to assess the efficiency of carrier-mediated ASO uptake by bacteria. Here, we have developed a “switch-on” reporter assay to measure ASO uptake efficiency in a semi-quantitative manner. The assay uses a synthetic RNA toehold switch fused to the mRNA of a fluorescent reporter protein, which is activated *in vivo* by a peptide nucleic acid (PNA)-based ASO upon delivery into the bacterial cytosol. We have used this assay to screen different cell penetrating peptides (CPPs) as ASO carriers in *Escherichia coli* and *Salmonella enterica* and observed up to 60-fold activation, depending on the CPP and bacterial strain used. Our assay shows high dynamic range and sensitivity, which should enable high-throughput screens for bacterial ASO carriers. We also show that the reporter can be used to study routes of PNA uptake, as demonstrated by reduced reporter activity in the absence of the inner membrane protein SbmA. In summary, we present a portable tool for the discovery of species-specific and efficient ASO carriers that will also be useful for a broader investigation of cellular uptake mechanisms of antibacterial ASOs.

**Importance:** The rise of antimicrobial resistance presents a major global health challenge. If not addressed, the death toll from resistant infections is expected to rise dramatically in the coming years. As a result, it is essential to explore alternative antimicrobial therapies. One promising approach is to target bacterial mRNAs using antisense oligomers (ASOs) to silence genes involved in essential functions, virulence, or resistance. However, delivering ASOs across bacterial membranes remains a major challenge and effective methods to monitor their uptake are limited. In this study, we develop a reporter assay to facilitate the high-throughput discovery of bacterial ASO carriers. This research paves the way for developing novel precision antisense-based antibacterial therapies.

## INTRODUCTION

Antimicrobial resistance is one of the major current global health challenges, emphasizing the need for novel strategies to treat resistant infections (1). Programmable species-specific RNA-based antibacterials are one promising alternative to conventional antibiotics (2–4). They are based on antisense oligomers (ASOs), i.e., short, single-stranded synthetic nucleic acid mimics designed to bind to complementary sequences in target mRNAs to repress their translation (5). By targeting mRNAs of essential genes, ASOs can lead to bacterial cell death (6). In addition, ASOs can be designed to suppress antibiotic resistance genes to reinstate the activity of antibiotics (7–11). Their use has been validated for multiple bacteria species, which include *Escherichia*, *Klebsiella*, *Pseudomonas*, *Salmonella,* and *Staphylococcus* species (12–17) and additional microbes (8, 18–20).

Currently, antibacterial ASOs are mostly based on peptide nucleic acid (PNA) or phosphorodiamidate morpholino (PMO) backbones due to their neutral charge, which is crucial for traversal of the negatively charged bacterial envelope (21). Nevertheless, ASOs alone are unable to translocate into the bacterial cytoplasm and need to be coupled to a carrier (2, 4). Common carriers include cell-penetrating peptides (CPPs), which are often short, cationic and amphiphilic peptides (22) that translocate across bacterial membranes using poorly understood mechanisms. While successful CPP-ASO conjugates have been reported for different bacterial species (8, 12–16, 18–20), our understanding of carrier properties required for optimal ASO delivery is limited.

To tackle this question, methods to assess the delivery potential of ASO carriers are required. Current methods either directly quantify CPP or ASO internalization or use indirect measurements of ASO-mediated gene regulation. To directly assess the uptake of the carrier module, CPPs can be labelled with fluorophores such as TAMRA (5-Carboxytetramethylrhodamine) to monitor their localization using flow cytometry followed by confocal microscopy (23, 24). Nevertheless, since the delivery potential of CPPs for cargos of different sizes and physicochemical properties varies (25) and ASOs are much larger than the TAMRA dye, this assay might not accurately measure PNA delivery. Instead, mass spectrometry has been used to directly detect ASOs in the bacterial cytoplasm or periplasm (21, 26). However, mass spectrometry-based methods are relatively laborious and limited in throughput. To assess ASO activity indirectly, bacterial growth inhibition can be used as a proxy if the ASO targets an essential gene. However, antibacterial activity is not a strict read-out for effective ASO delivery, because the CPP moiety itself may also be toxic (12, 13, 23). Monitoring phenotypic changes post CPP-ASO treatment is another option to evaluate the delivery potential of carriers. For example, inhibition of the *ftsZ* gene encoding an important cell division protein causes bacterial filamentation (27), which can easily be scored under the microscope. Alternatively, targeting the mRNAs of reporter proteins such as beta-galactosidase (encoded by the *lacZ* gene) (28), luciferase (29), or fluorescence proteins (30) can be used. These reporter assays have the advantage of being non-destructive, but loss of gene expression can also result from non-specific toxicity of the CPP-ASO conjugates.

Here, we have developed a translational “switch on” reporter assay to assess ASO delivery into the bacterial cytoplasm. It draws inspiration from a common post-transcriptional mechanism of gene activation by small regulatory RNAs (sRNAs) in bacteria, i.e., sRNA-mediated disruption of an inhibitory RNA structure around the ribosome binding site (RBS) of a target mRNA (31). To design our reporter we chose an established synthetic RNA toehold switch (32), referred to as TS7, that sequesters the RBS and start codon of the mRNA of a fluorescent protein in a stem-loop structure (Fig. 1). The switch can be activated by co-expression of antisense RNAs that bind the 5’ flank of the stem-loop (32). We designed PNA-based ASOs that similarly bind the switch and activate the synthesis of the fluorescent reporter protein, resulting in an “on-state”. Variation in fluorescence resulting from this “on-state” can be used to infer the efficacy of CPPs to deliver ASOs into bacteria, independent of their antibacterial activity. We tested this reporter in different bacterial strains using variable ASO concentrations, growth media and fluorescent reporter proteins. We also successfully conducted a small screen of 10 CPPs. This sets the stage for high-throughput screens for new ASO carriers, be these short peptides or other delivery vehicles for transversal of the bacterial envelope.

**Figure 1.**
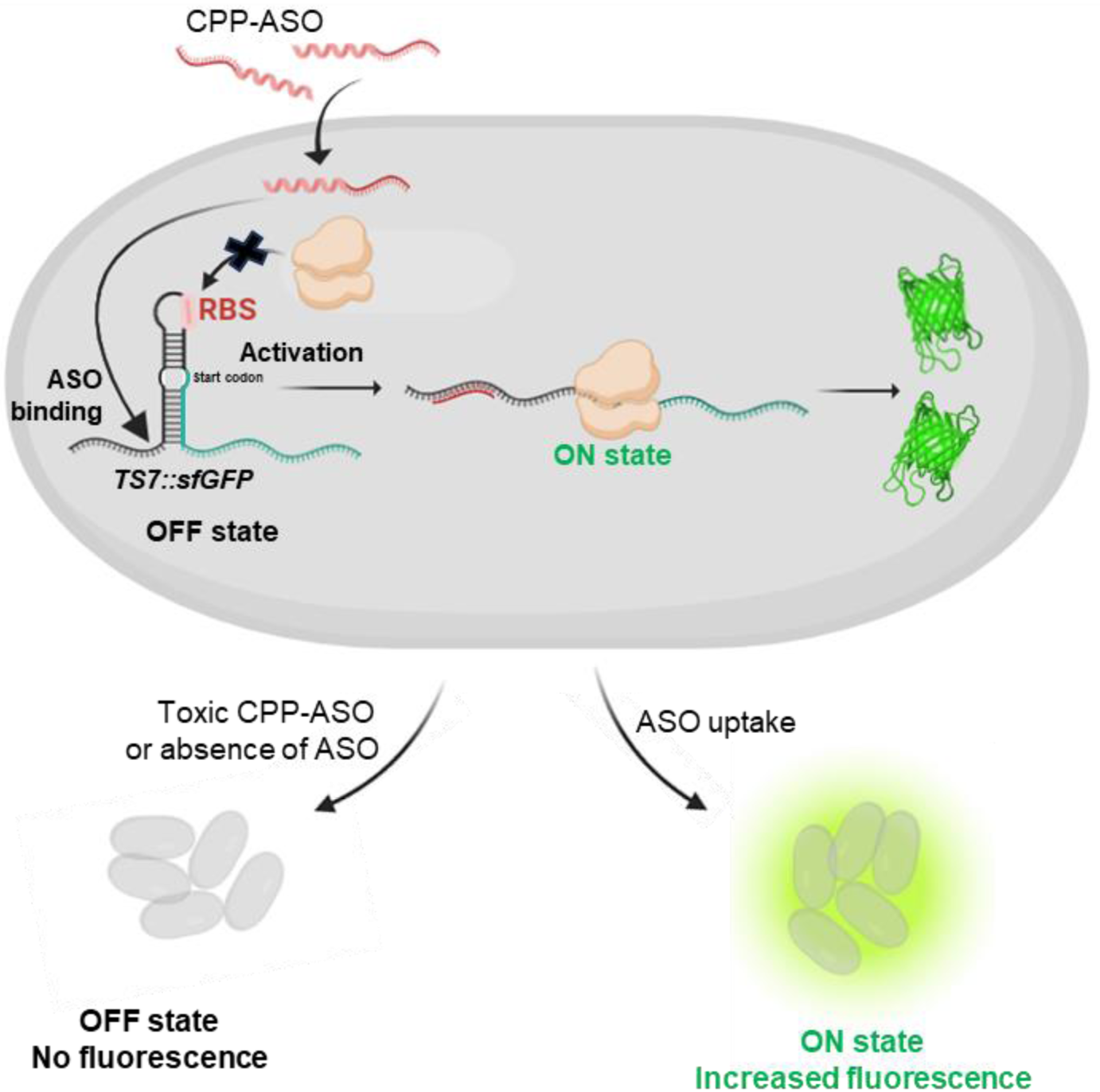
Design rationale for the antisense activation-based “switch-on” reporter assay. Schematic overview of antisense oligonucleotide (ASO)-based RNA toehold switch activation assay. In the OFF state, the TS7 toehold switch sequesters the ribosome binding site (RBS) and the start codon, thus preventing protein synthesis of the downstream reporter gene encoding superfolder GFP (*sfGFP*). Upon antisense uptake, the ASO hybridizes with the stem-loop of TS7 and opens the inhibitory secondary structure. The RBS becomes accessible and reporter protein translation is initiated (ON state). In case of inefficient CPP-PNA delivery, the reporter will stay in its off-state.

## RESULTS

### PNA-mediated activation of the RNA toehold switch in vitro

We build the ASO sensor component of our switch-on reporter based on the TS7 toehold switch, an 83 bp sequence, which, when transcribed, can form an imperfect 18 bp long RNA hairpin (Fig. 2A), as reported by Green *et al*. (32). In the original description, TS7 was activated by the co-expression of a *trans*-acting antisense RNA with 30 nucleotides complementarity to the 5’ untranslated region of the toehold switch (32). We hypothesized that due to the higher affinity of PNA to RNA (33), shorter PNAs in the typical length range of antibacterial ASOs could achieve a similar effect. To test if 11mer PNAs can activate the toehold switch, we first performed an *in vitro* translation assay. We titrated *in vitro* transcribed *TS7*::*gfp* RNA with varying concentrations of a PNA that targets the toehold region of the TS7 switch (PNA_toe) (Fig. 2A). Accumulation of GFP after *in vitro* translation was detected by western blot (Fig. 2B). The addition of PNA_toe led to a dose-dependent increase in GFP levels, reaching ∼11-fold upregulation at 10x molar excess of PNA over RNA compared to the water control (Fig. 2C). A non-targeting PNA (PNA_ctrl) had no effect even when used at a 10x molar excess over *TS7::gfp* RNA. This *in vitro* assay demonstrates that the 11mer PNA_toe can activate the TS7 RNA toehold switch for the synthesis of the fused reporter protein.

**Figure 2.**
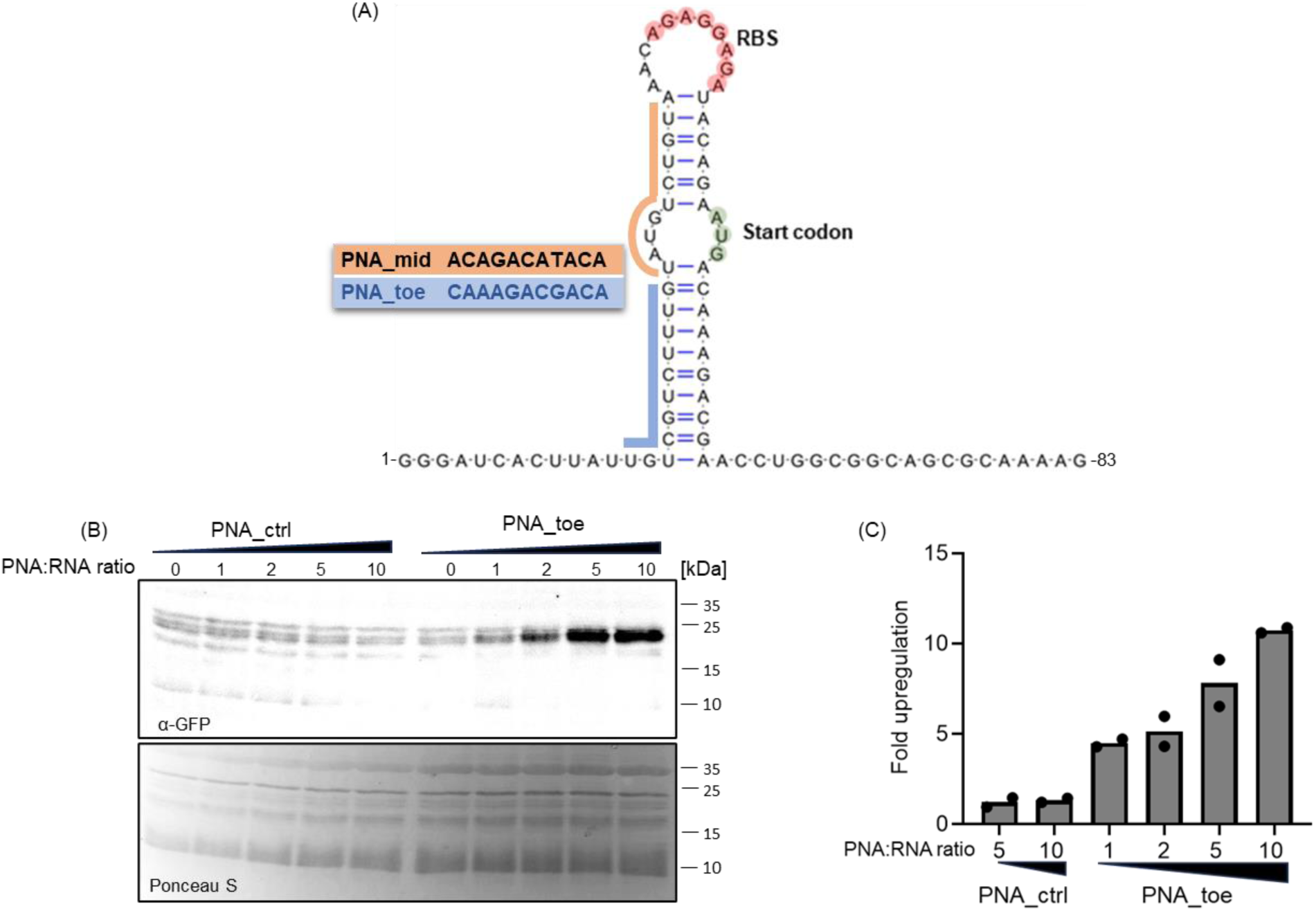
Antisense-mediated activation of toehold switch-controlled GFP expression *in vitro*. (A) Structure of the TS7 toehold switch. The RBS and translational start codon are highlighted in red and green, respectively. The PNA sequences with their respective target sites are indicated in blue and orange. (B-C) *In vitro* transcribed *TS7::gfp* RNA was subjected to an *in vitro* translation assay. Synthetic *TS7::gfp* fusion mRNA was incubated with the indicated molar excesses of a non-targeting PNA_ctrl or PNA_toe. As a negative control, an equal volume of water was added to the samples (“0”). RNA was translated *in vitro* and samples were separated on 12% SDS-PAA gels with subsequent blotting on nitrocellulose membranes. (B) Western blot-based detection of GFP was performed using a monoclonal α-GFP antibody (upper panel). PonceauS staining was used to control for equal loading (lower panel). (C) Changes in GFP protein levels were quantified using ImageJ. Relative protein expression levels of GFP were calculated based on the water control sample. Bars represent the mean of two independent experiments, individual data points are shown as well.

### Activation of the toehold switch upon PNA delivery in vivo

Next, we tested whether 11mer PNAs can activate the toehold switch in bacteria. In constructing a reporter plasmid, we cloned the TS7 sequence upstream of the coding sequence of superfolder GFP (sfGFP) and downstream of a constitutive P_LtetO-1_ promoter in the pXG10-SF vector (34) (see Methods for details). The final product is a fusion protein (TS7::sfGFP) in which the ten amino acids from the short TS7 coding sequence are linked to the N-terminus of sfGFP. We introduced the reporter plasmid in the model organism *Salmonella enterica* serovar Typhimurium, referred to as *Salmonella* hereafter. The bacteria were grown to an early log-phase in standard Mueller-Hinton broth (MHB), commonly used for testing antibacterial activity (35). As a proof-of-concept, we chose to deliver the PNAs through conjugation with (KFF)_3_K, a frequently used CPP (referred to as KFF hereafter). We tested two PNAs, PNA_toe as used above *in vitro* and a PNA that targets the mid-stem region (PNA_mid) of the switch (Fig. 2A). We treated the bacterial culture with varying concentrations (2.5 µM, 5 µM or 7.5 µM) of the KFF-PNAs in a microtiter plate, recording GFP fluorescence and turbidity as bacterial growth continued. We did not use higher concentrations to avoid unspecific side effects. While treatment with PNA_ctrl or water caused no substantial increase in fluorescence over time, both PNA_toe and PNA_mid led to a concentration-dependent increase in fluorescence, with PNA_toe being more effective (Fig. 3A and S1). Moreover, PNA_toe had no effect on bacterial growth, whereas PNA_mid impaired bacterial growth (Figs. 3A). Based on both, its superior ability to activate the reporter and its lower toxicity, we selected PNA_toe as the activating ASO moiety for further experiments.

**Figure 3.**
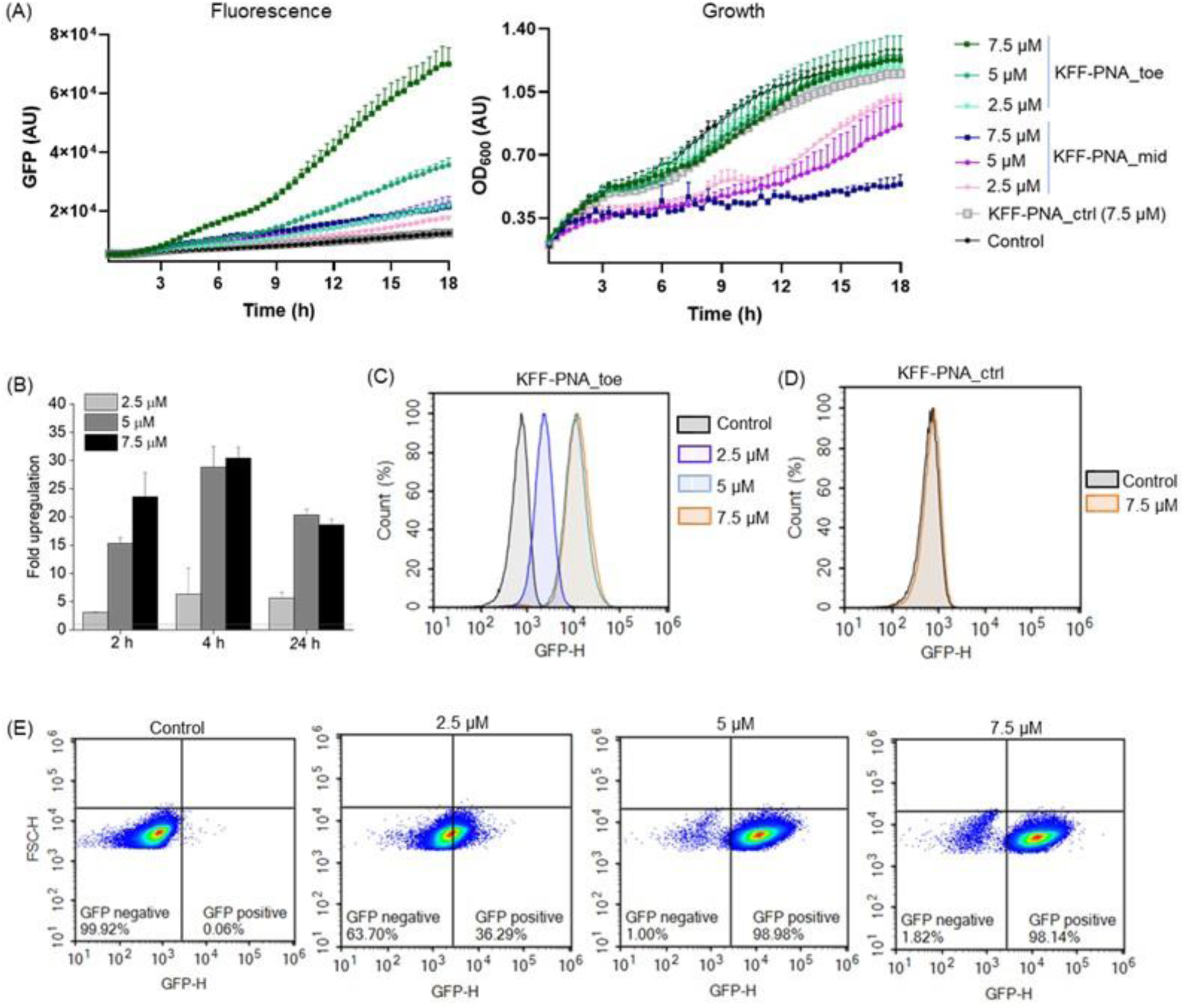
KFF-PNA mediated activation of toehold switch-controlled sfGFP expression in *Salmonella*. *Salmonella* carrying the *TS7::sfgfp* encoding plasmid were treated with increasing concentrations of KFF-PNA_toe or KFF-PNA_mid. As negative controls, a non-targeting PNA (KFF-PNA_ctrl) or water (Control) were used. (A) Fluorescence (sfGFP) (left) and bacterial growth (right) were recorded in a microplate reader for 18 h post-PNA treatment. Data points indicate mean sfGFP fluorescence intensities (left panel) and the optical densities at 600 nm (OD_600_; right panel). Error bars show the standard deviation of the mean fluorescence and OD from two replicates. (B-E) Flow cytometric analysis of sfGFP induction post PNA treatment. (B) Concentration-dependent sfGFP induction after treatment with KFF-PNA_toe for 2 h, 4 h, and 24 h. Bars indicate the relative upregulation of PNA-treated samples compared to the water-treated control. Upregulation was calculated based on the median fluorescence intensities. Error bars indicate the standard deviation of median fluorescence intensity from two independent experiments. (C, D) Flow cytometry histograms showing changes in sfGFP fluorescence 4 h post-treatment with (C) KFF-PNA_toe at 7.5 µM, 5 µM and 2.5 µM, or (D) a non-targeting PNA (KFF-PNA_ctrl) at 7.5 µM. (E) Density plots of sfGFP induction in bacterial populations post-treatment with KFF-PNA_toe at the indicated concentrations. For the control (left panel), an equal volume of water was added. AU indicates arbitrary units. Each experiment was performed at least two times.

In contrast to plate reader-based measurements, which provide values of the population average, flow cytometry offers a more quantitative readout with single-cell measurements (36). Using flow cytometry, we validated the observations from the microplate reader, confirming both the concentration-dependent and time-dependent variation in GFP fluorescence after PNA treatment (Figs. 3B-C and S2). Interestingly, a decrease in the fluorescence was observed at 24 h relative to 4 h post-treatment, potentially due to degradation of the KFF moiety in the medium (26), which renders the remaining PNA constructs unable to enter the bacterial cells. As expected, PNA_ctrl did not activate GFP expression (Fig. 3D). The flow cytometry density plots revealed a gradual homogenous increase in the percentage of the GFP positive population with increasing PNA concentration (Fig. 3E). The dynamic range of the flow cytometry readout (determined as the highest upregulation observed with the KFF-PNA compared to water-treated control) exceed that of the microplate reader (1.5-30 *vs* 1.5-10, respectively). Since treatment with either 7.5 µM or 5 μM KFF-PNA_toe showed equally strong activation of the reporter 4 h post-treatment, we settled on 5 μM for all further experiments to minimize potential toxicity.

### An improved toehold switch enhances PNA-mediated activation

Our activation assay relies on the ability of an ASO to either open or prevent the inhibitory stem-loop of the RNA toehold switch from forming, for mRNA translation to initiate. We reasoned that we might be able to improve the dynamic range and sensitivity of the assay by weakening the stem-loop for easier melting. To this end, we mutated positions 9, 13, and 18 of the stem, all of which are outside the loop and binding site of PNA_toe (Fig. 4A). These mutations caused a slight increase in basal reporter protein synthesis as seen by higher fluorescence without PNA treatment (Fig. 4B). Seeking the highest signal-to-background ratio post-PNA treatment, we treated cells carrying these mutated switch variants with KFF-PNA_toe for 4 h. While KFF-PNA_toe resulted in around 30-fold upregulation of the reporter with the original sequence (Fig. 3B), we observed a ∼60-fold activation of the switch mutated at position 9 (TS7-Mut_9, Fig. 4C). By contrast, mutations at positions 13 (TS7-Mut_13) or 18 (TS7-Mut_18) led to no improvement over the original switch. We also examined fluorescence 2 h after treatment to assess if the mutations facilitated faster activation, but this was not the case (Fig. 4C). Since TS7-Mut_9 offered the highest dynamic range, we selected this reporter for the experiments below.

**Figure 4.**
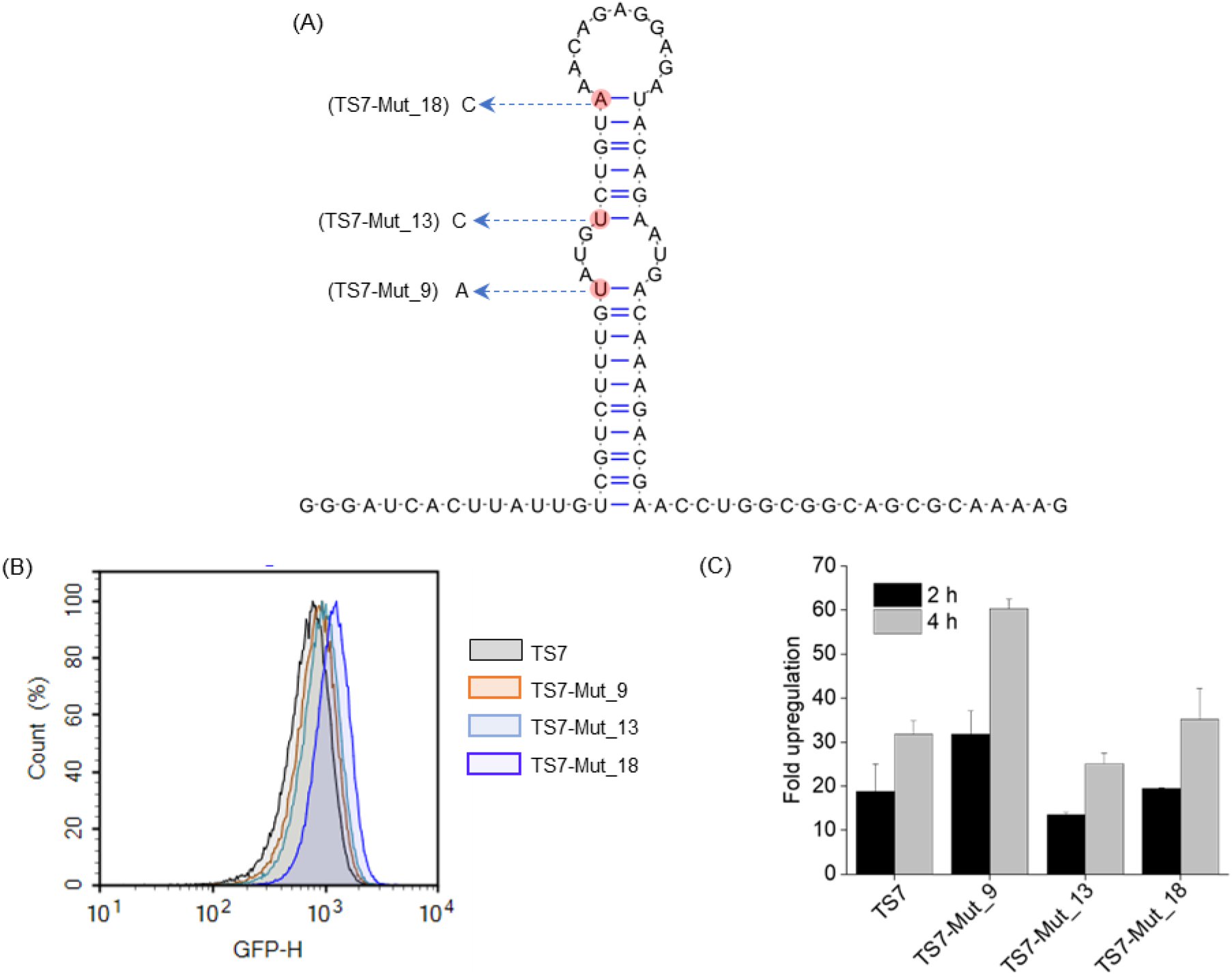
Impact of single-nucleotide mutations in the stem of the toehold switch (TS7) on the reporter’s sensitivity. (A) Point mutations (indicated in red) introduced within the *TS7::sfGFP* toehold switch to improve response to PNA-mediated activation. (B, C) Flow cytometric analysis of *Salmonella* bearing plasmids containing the original or mutated *TS7::sfGFP* reporter constructs. (B) Comparative representation of basal fluorescence intensities of mutated and original toehold switch constructs measured by flow cytometry. (C) Bacterial cells containing the different *TS7::sfGFP* constructs were treated with 5 µM of KFF-PNA_toe and sfGFP signals were measured at 2 h and 4 h post-treatment. Bars indicate relative upregulation of sfGFP calculated based on the median fluorescence intensities of PNA-treated samples relative to the water control. Error bars indicate the standard deviation calculated from two independent experiments.

### Influence of growth medium and fluorescent proteins on reporter activation

The composition of the culture media can influence the interaction between cationic CPPs and bacterial cells (37, 38) and so might affect CPP-mediated PNA delivery. To study how different media affected CPP-PNA-mediated reporter activation, we compared the upregulation of GFP expression after 4 h treatment in three different standard culture media: MHB (used above), a minimal medium referred to as M9 and nutrient-rich Luria Bertani broth (LB). In both MHB and M9, KFF-PNA_toe triggered a ∼60-fold activation of the reporter (Fig. 5A-B). In contrast, we observed only ∼16-fold upregulation in LB. This shows that the activity of the reporter assay depends on the culture medium. Given that MHB is commonly used for testing antimicrobials, we continued with MHB for all subsequent studies.

**Figure 5.**
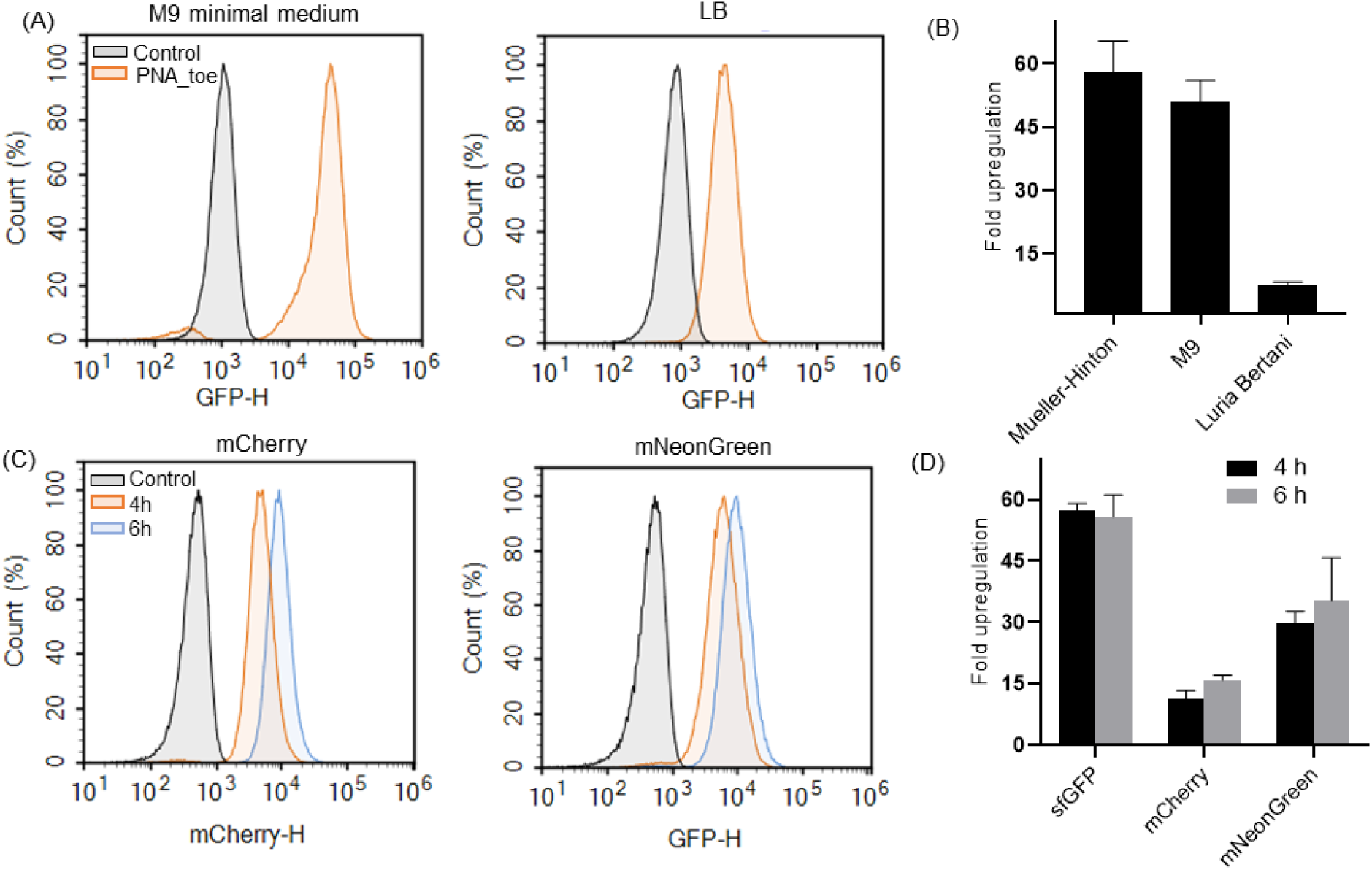
Effect of culture media and fluorescent reporters on the degree of PNA-mediated toehold switch activation in S*almonella*. (A) Flow cytometry histograms of sfGFP fluorescence of *Salmonella* containing the *TS7::sfGFP-Mut_9* reporter construct treated with KFF-PNA_toe at 5 µM in M9 minimal medium (left) or Luria-Bertani (LB) broth (right) for 4 h. Control indicates bacteria treated with an equal volume of water. (B) Fold upregulation of sfGFP fluorescence in different culture media at 4 h post-treatment relative to water-treated control cells. (C) Flow-cytometry histograms of fluorescence of *Salmonella* containing the *TS7-Mut_9::mCherry* (left) or *TS7-Mut_9::mNeonGreen* (right) reporter constructs, treated with KFF-PNA_toe at 5 µM in Mueller Hinton medium for 4 h and 6 h. (D) Fold upregulation of mCherry or mNeonGreen fluorescence post-treatment with KFF-PNA_toe at 5 μM. (B, D) Fold upregulation of fluorescence was calculated based on the median fluorescence intensities of PNA-treated samples relative to the water-treated control, measured by flow cytometry. Error bars indicate the standard deviation calculated from two independent experiments.

To assess how the reporter protein influences the dynamic range of the assay, we tested two additional commonly used fluorescent proteins, mNeonGreen and mCherry. In accordance with the fact that sfGFP is among the brightest fluorescent proteins and characterized by a rapid maturation time in Enterobacteriaceae (39), the sfGFP-based reporter showed the highest upregulation after 4 h and 6 h (Fig. 5D). Both mCherry and mNeonGreen showed a lower fold activation than sfGFP at both time points, potentially due to longer maturation times and poor stability, at least for mCherry (40). Since sfGFP showed higher and faster upregulation than both mCherry and mNeonGreen, we continued with the sfGFP reporter.

### The toehold switch reporter can provide insights into CPP-PNA uptake mechanisms

To confirm that activation of the toehold-switch reporter correlates with successful uptake of PNAs, we used a *Salmonella* Δ*sbmA* mutant strain unable to express the inner membrane transporter SbmA. This protein has been shown to mediate the uptake of KFF-conjugated PNAs into the taxonomically close relative, *E. coli* (27). In the Δ*sbmA* strain, treatment with KFF-PNA_toe triggered only marginal GFP expression in a small subpopulation of cells (ca 20 %) as opposed to nearly 100 % of all cells in the corresponding wild-type strain (Fig. 6A). The overall fluorescence intensity of the Δs*bmA* mutant was also much lower compared to the wild-type (Fig. 6B). These results validate that the assay reports PNA uptake and indicate that the reporter can also be used to assess the involvement of bacterial transporters in PNA delivery.

**Figure 6.**
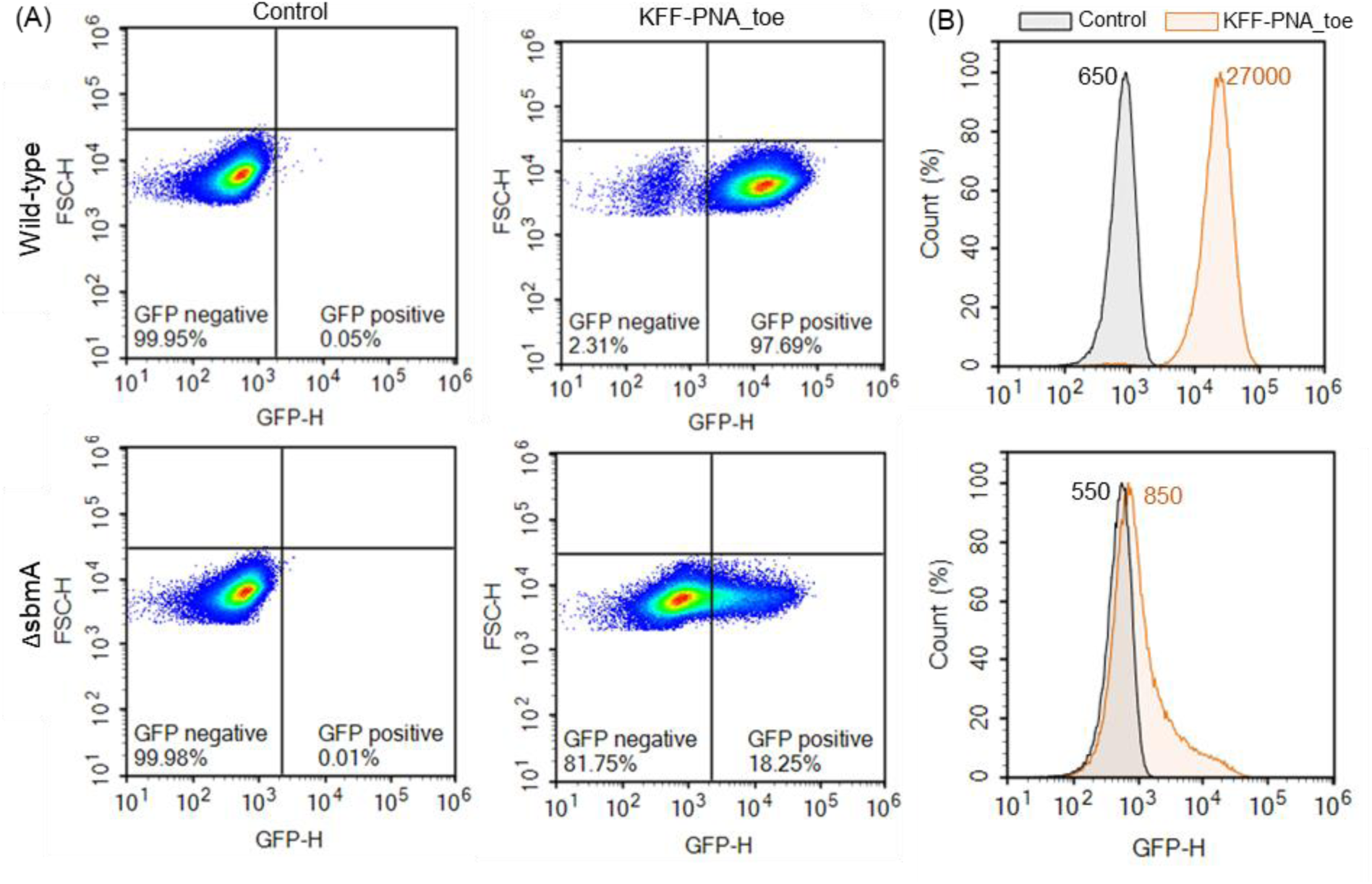
Flow cytometric analysis of CPP-PNA uptake using the toehold switch reporter in a Δ*sbmA* mutant of *Salmonella*. Flow cytometric (A) density plots and (B) histograms depicting the heterogeneity in sfGFP expression upon activation of *TS7-Mut_9::sfGFP* 4 h post-treatment with KFF-PNA_toe (5 µM). The upper panels show results for *Salmonella* wild-type, and the lower panels depict the results for an isogenic Δ*sbmA* mutant. Control indicates bacteria treated with an equal volume of water. (B) Median fluorescence intensity values for water-treated control (Control, black) and PNA-treated (KFF-PNA_toe, orange) samples are given in the plots.

### The toehold switch reporter enables the assessment of CPP-mediated PNA delivery

Having established that the reporter assay reads out PNA delivery, we applied it to assess new CPP candidates. We tested a small library of 10 CPPs for their ability to deliver PNA_toe into *Salmonella*. We included the established CPPs KFF, RXR, TAT, and ANT, all of which are known to deliver PNA into *Salmonella* (13). We also included six other CPPs with varying cationicity and length with reported applications in prokaryotic or eukaryotic cells (24, 41) (Table 1). To rule out non-specific activation of the reporter, CPPs conjugated to the non-targeting PNA_ctrl were tested; none of these conjugates activated the reporter (Table S1). In addition to fluorescence, we simultaneously measured bacterial growth, with potential growth impairment flagging toxicity of the CPP moiety. Although there were no strong toxic effects, we did observe mild growth retardation for RXR and Seq23 (Fig. 7A). To control for these effects, fluorescence was normalized to the optical density of the bacterial culture (Fig. S3). We observed a rapid increase in fluorescence signal for the PNA_toe conjugates of KFF and RXR, while the other CPPs showed a slower increase (Fig. 7B). At 17 h post-treatment, KFF, RXR, and ANT exhibited the highest fold increase in fluorescence compared to the water-treated control, followed by R5, Seq118, and Seq35 (Fig. 7C). A negligible upregulation was observed with TAT, R9-TAT, and L5a, and none with Seq23. These data demonstrate that our reporter assay can be used to screen CPP candidates by using the microplate reader.

**Figure 7.**
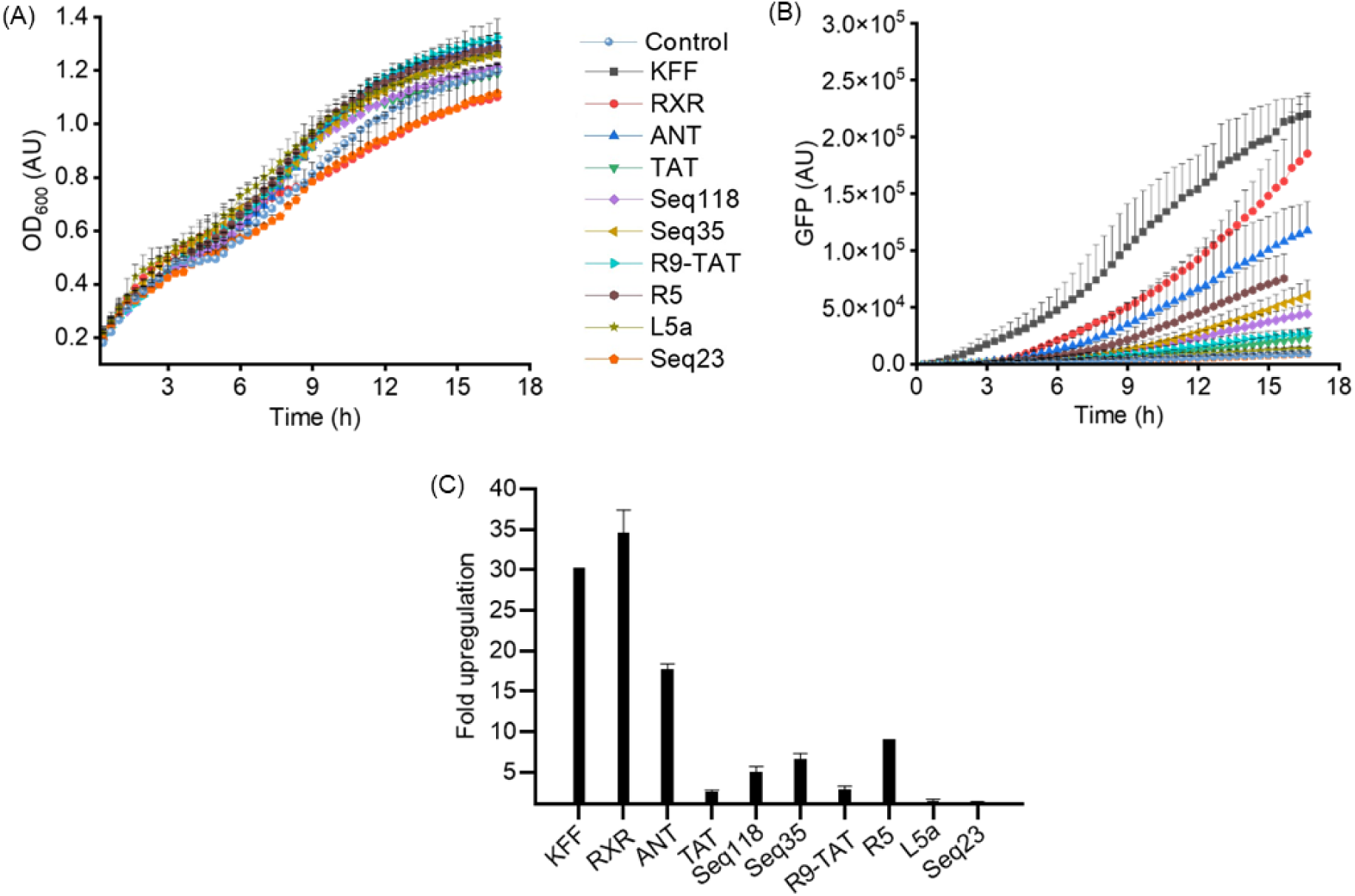
Antisense activation of toehold switch reporter enables screening of CPPs for PNA delivery. (A-C) *Salmonella* carrying the *TS7-Mut_9::sfGFP* encoding plasmid were treated with different CPP-PNA_toe constructs at 5 µM. As negative control, an equal volume of water (Control) was added instead. (A) Bacterial growth (OD_600_) and (B) sfGFP fluorescence (GFP) were recorded in a microplate reader for 17 h post-PNA treatment. AU, arbitrary units. (C) Relative upregulation of sfGFP fluorescence was calculated based on the fluorescence intensities at 17 h of PNA-treated samples relative to the water control. Error bars indicate the standard deviation of fold upregulation calculated from two independent experiments.

**Table 1.** Description of cell-penetrating peptides (CPP) used in this study.

| CPP | Sequence * | Number of |  |
| --- | --- | --- | --- |
|  |  | Cationic amino acids | Hydrophobic groups |
| KFF | KFFKFFKFFK | 4 | 6 |
| RXR | RXRRXRRXRRXRB | 8 | 6 |
| ANT | RQIKIWFQNRRMKWKK | 7 | 6 |
| Tat | GRKKRRQRRRYK | 9 | 1 |
| R9-Tat | GRRRRRRRRRPPQ | 9 | 0 |
| Seq118 | RRRQRRKKRGY | 8 | 1 |
| Seq35 | RKKRRQRR | 7 | 0 |
| R5 | RRRRR | 5 | 0 |
| L5a | RRWQW | 2 | 2 |
| Seq23 | VALLPAVLLA | 0 | 9 |

\* CPP sequences are shown from N to C termini

Having established the CPP screening assay in *Salmonella*, we next tested it in *E. coli*. We chose the *E. coli* K12 strains BW25113 and MG1655 and the uropathogenic isolate *E. coli* 536 (UPEC 536). Using the same reporter (*TS7-Mut_9::sfGFP*) and conditions as above, we observed the highest reporter activation for KFF and RXR conjugates in all these three *E. coli* strains, similar to *Salmonella* (Fig. 8). Seq23 remained the least effective PNA carrier. Importantly, *E. coli* BW25113 and MG1655 displayed ∼10-fold lower fluorescence signals for all CPPs compared to UPEC despite similar basal fluorescence in all strains (Fig. 8). For these strains, a more sensitive method compared to the microplate reader such as flow cytometry might provide better results.

**Figure 8.**
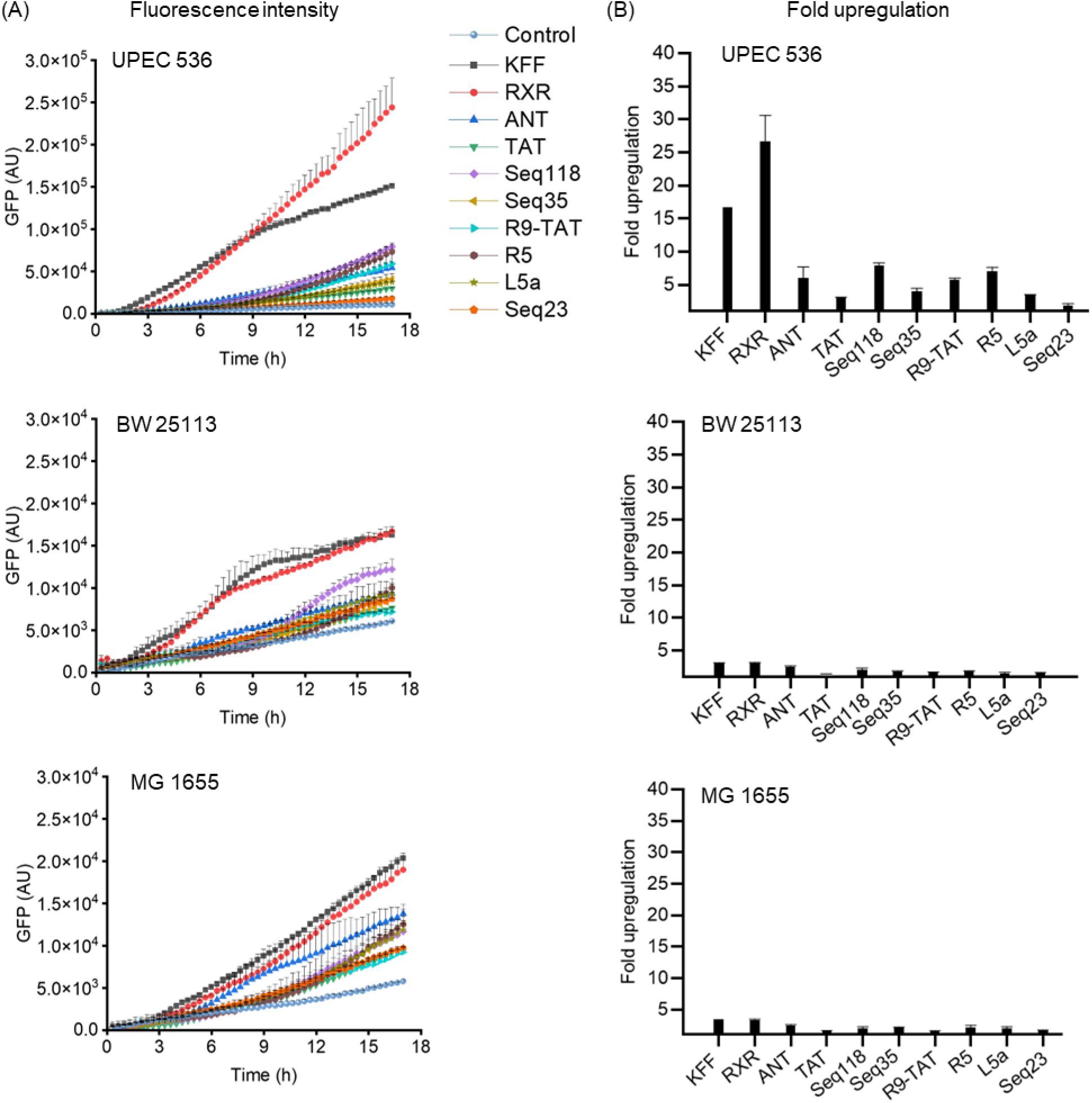
Examining the performance of the toehold switch assay for screening of CPPs against *E. coli*. Various *E. coli* strains carrying the *TS7-Mut_9::sfGFP* encoding plasmid were treated with different CPP-PNA_toe constructs at 5 µM. As negative control, an equal volume of water (Control) was added instead. (A) Fluorescence intensity of sfGFP was recorded in a microplate reader for 17 h post-treatment measured in a microplate reader. (B) Upregulation in sfGFP fluorescence calculated relative to the water control based on the normalized fluorescence intensities at 17 h post-treatment. The experiment was performed two times and bars indicate the mean value; error bars indicate the standard deviation from two independent experiments.

### Flow cytometry shows dynamic differences in toehold switch activation by CPP-PNAs

According to previous RNA-seq analyses (12, 13, 21), CPP-PNAs can strongly downregulate target mRNAs within 15 min. This suggests that reporter activation could possibly be scored earlier, permitting more sensitive detection of GFP fluorescence. We therefore repeated the screen and assayed fluorescence by flow cytometry at 4, 6, and 17 h post-treatment. In all strains, KFF, RXR, and ANT showed rapid upregulation at 4 h, whereas Seq35, Seq118, R9-TAT, and R5 only showed upregulation at the 17 h time point (Fig. 9); Seq23 consistently was the least active carrier. The level of upregulation observed in the *E. coli* K12 strains was again substantially lower than in *Salmonella* and UPEC. Nevertheless, as compared to the plate reader, flow cytometry showed a higher dynamic range and sensitivity to study the efficacy of CPP-mediated ASO delivery in all the tested bacteria (Fig. 10). Interestingly, we observe CPP-dependent differences in the dynamics of ASO-mediated sfGFP induction. For example, while fluorescence increases over time for most CPPs, KFF showed a decrease (Fig. 9). This might be related to the stability of KFF in the culture media. Overall, our results show that CPP-mediated PNA delivery varies over time and in different bacterial strains.

**Figure 9.**
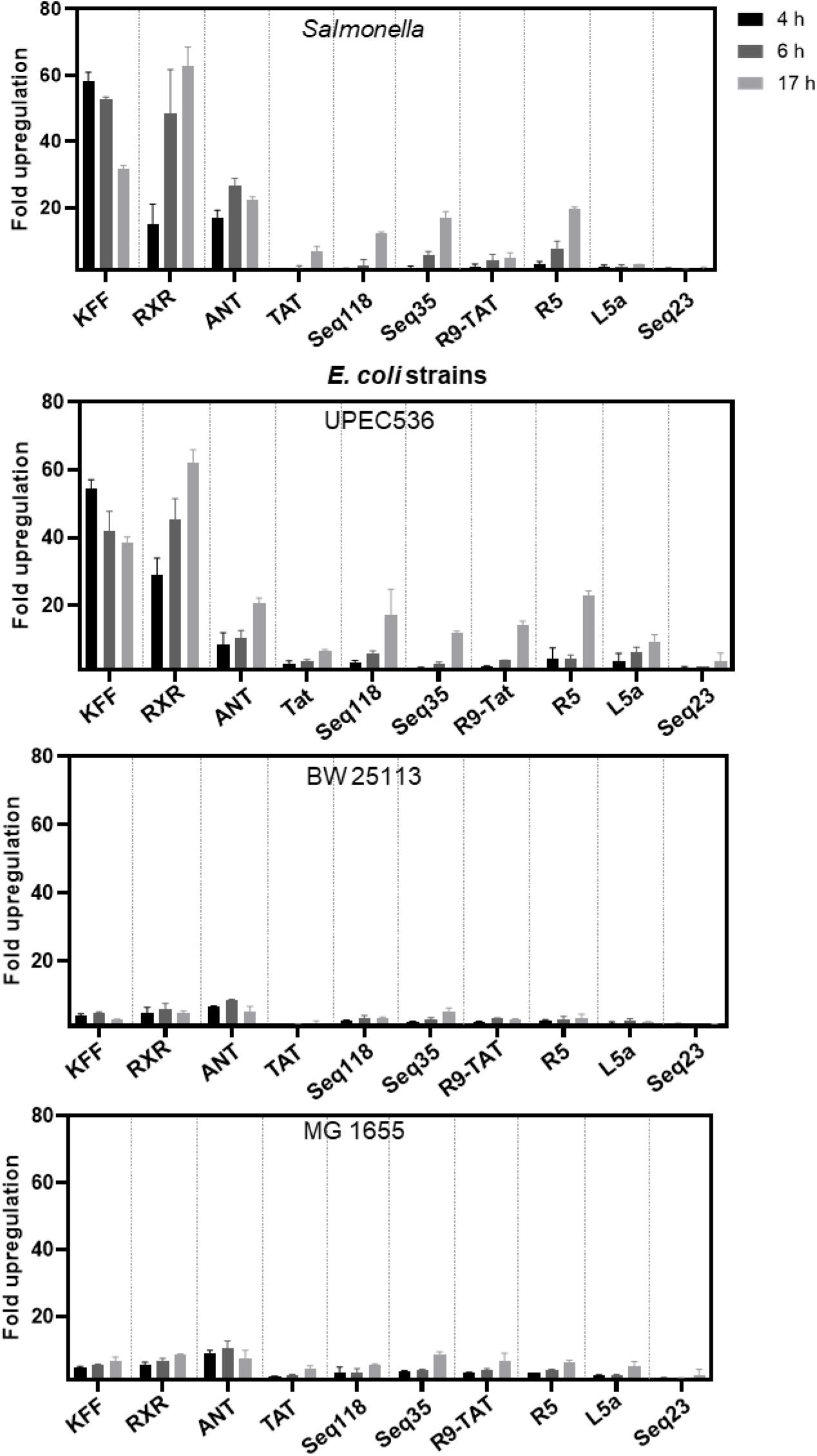
Flow cytometric analysis of fluorescence of sfGFP informs on delivery efficiency of CPPs. Bacterial cultures expressing the *TS7-Mut_9::sfGFP* reporter construct treated with the indicated CPP-PNA_toe constructs at 5 µM each. The median fluorescence intensity of sfGFP was measured after 4 h (black bars), 6 h (dark grey bars) and 17 h (light grey bars) of treatment using flow cytometry. Relative upregulation of sfGFP is shown and was calculated by dividing the median fluorescence intensity of the test samples by the median fluorescence intensity of the control samples, to which water was added. Error bars indicate the standard deviation calculated from two independent experiments.

**Figure 10.**
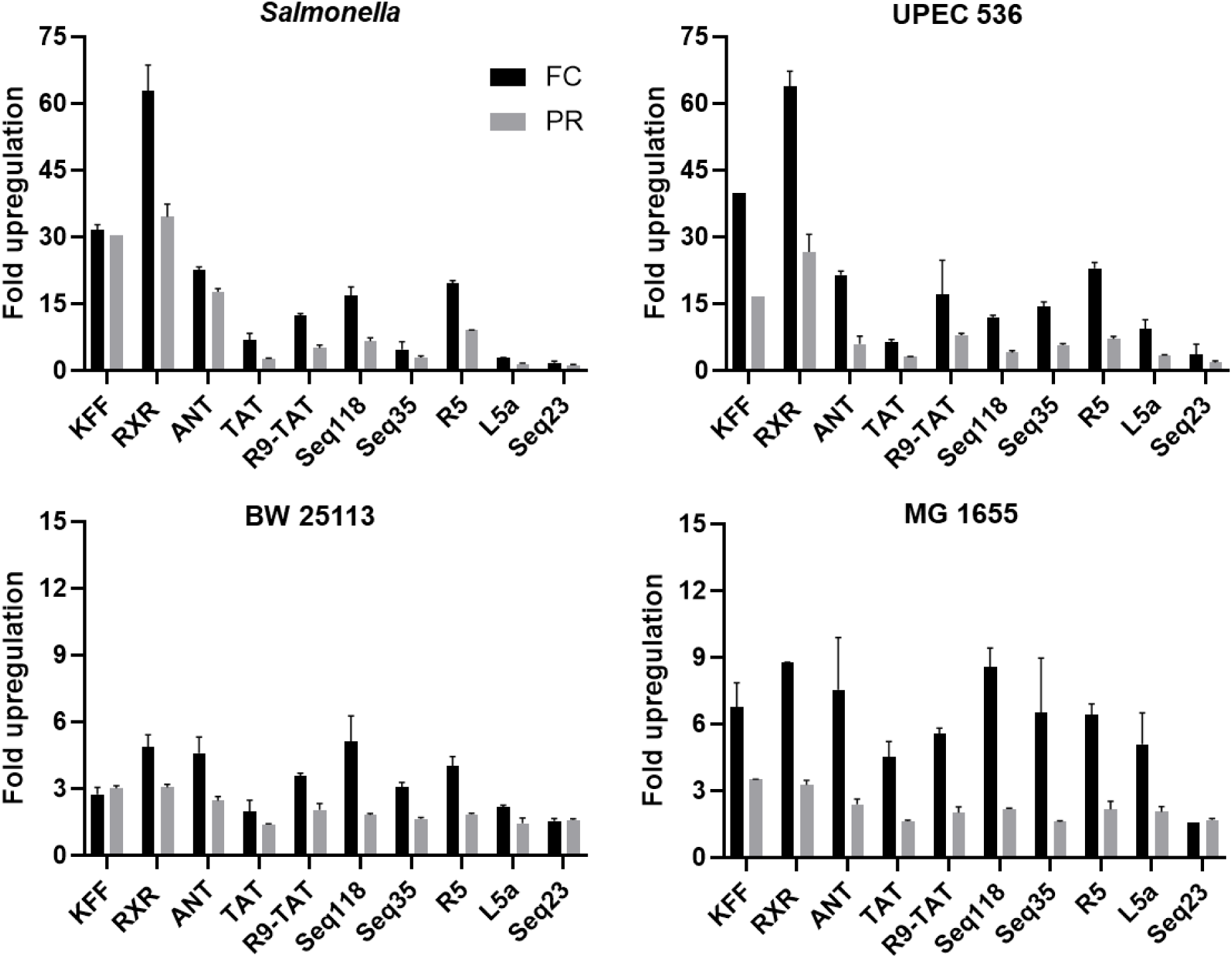
Comparison of CPP-PNA_toe-mediated reporter activation evaluated by flow cytometry (FC) *versus* microplate reader (PR). Average fold upregulation in sfGFP fluorescence was calculated relative to the respective water-treated controls. Fluorescence was measured 17 h post-treatment with various CPP-PNA_toe constructs using microplate reader (grey bars; see also Figure 8) or flow cytometry (black bars; see also Figure 9). Error bars indicate the standard deviation of fold upregulation from two independent experiments.

### Reporter activation correlates with the activity of antibacterial CPP-PNA conjugates

To assess if reporter activation correlates with antibacterial activity, we determined the minimum inhibitory concentration (MIC) of the CPPs tested above when conjugated to the antibacterial 10mer PNA sequence that targets the mRNA of *acpP* (acyl carrier protein), a well-known target of antibacterial PNAs (13, 42). We observed that RXR, KFF, and ANT, which showed rapid and marked upregulation in the reporter assay also showed low MICs (<2.5 μM against *Salmonella* and UPEC; <5 μM against the K12 strains) (Fig. 11 and S4, Table 2). The Seq23 CPP-PNA conjugate, which showed no reporter activation, had no antibacterial activity (MIC >10 μM). In general, the two *E. coli* K12 strains (BW25113, MG1655) were less susceptible to CPP-*acpP* treatment than *Salmonella* and UPEC, again consistent with the reporter assays (Fig. 11). We also observed discrepancies between reporter activation and antibacterial activity. Seq118, which activated the reporter in *E. coli* BW25113 by 5-fold, failed to inhibit bacterial growth of this strain. L5a barely activated the reporter in *Salmonella* (<3-fold) but mediated strong inhibition of both *Salmonella* and UPEC (MIC ≤ 2.5 µM) (Fig. 11). Despite these inconsistencies, results of the switch-on assay generally correlate with the antibacterial assay, demonstrating potential of our reporter for identifying new carriers for antisense antibiotics.

**Figure 11.**
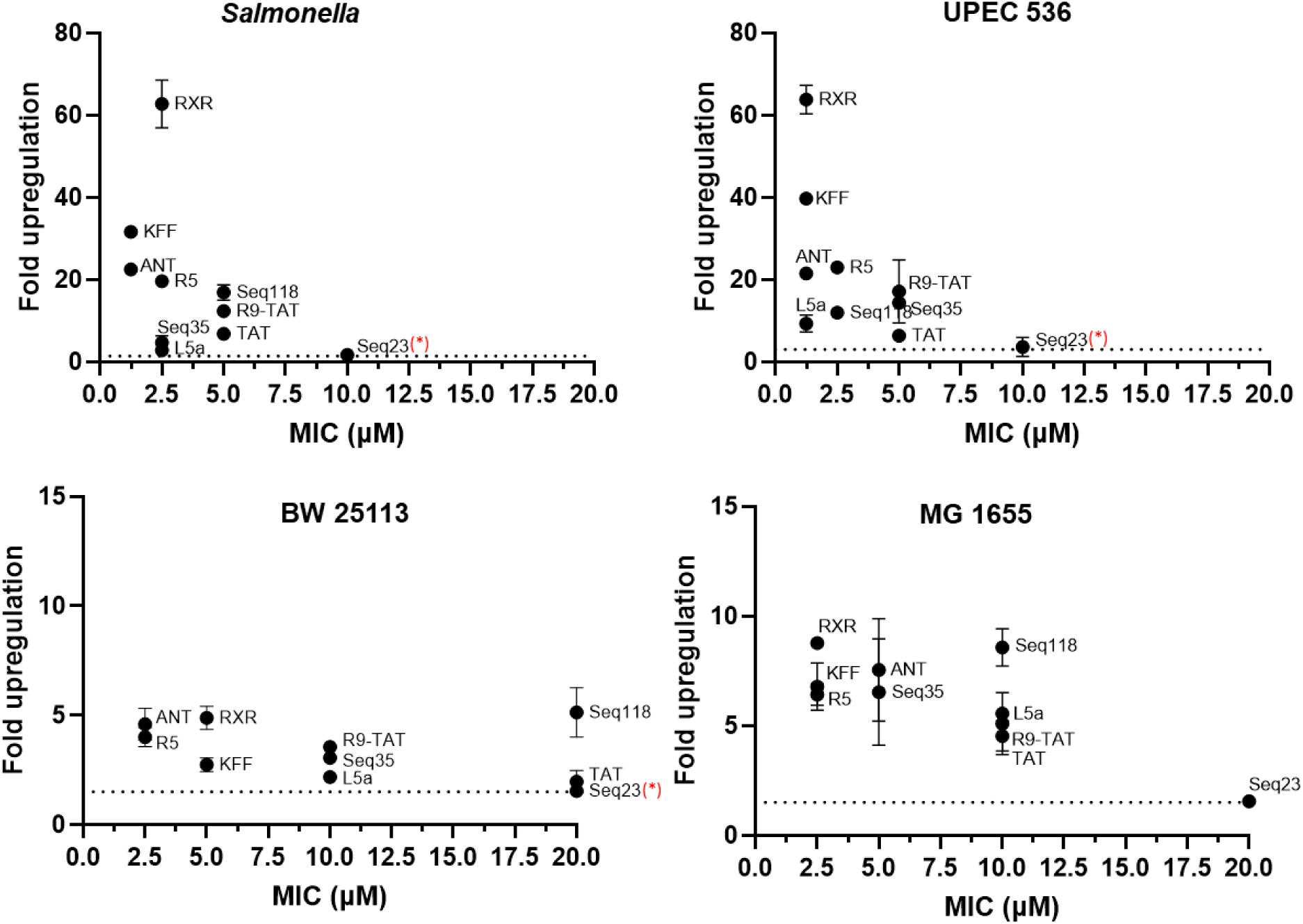
Comparison of CPP-PNA_toe-mediated activation with the growth inhibitory capacity of CPP-PNA constructs targeting the essential gene *acpP*. CPP-PNA-mediated activation of the reporter assay was plotted against the growth inhibitory activity of CPP-*acpP* constructs. Average fold upregulation in sfGFP fluorescence (y-axis) was calculated relative to the respective water-treated controls. Fluorescence was measured 17h post-treatment with various CPP-PNA_toe constructs using flow cytometry (shown in Fig. 9). Cognate CPP-PNA constructs targeting the mRNA of the essential gene *acpP* were used to determine their minimal inhibitory concentration (MIC) (x-axis). Each data point indicates the mean value of sfGFP induction and MIC, and error bars indicate the standard deviation of two independent experiments. ‘*’ indicates the MIC could not be determined for Seq 23, and is >10 μM in *Salmonella* and UPEC, >20 μM in the *E. coli* BW25113.

**Table 2.**
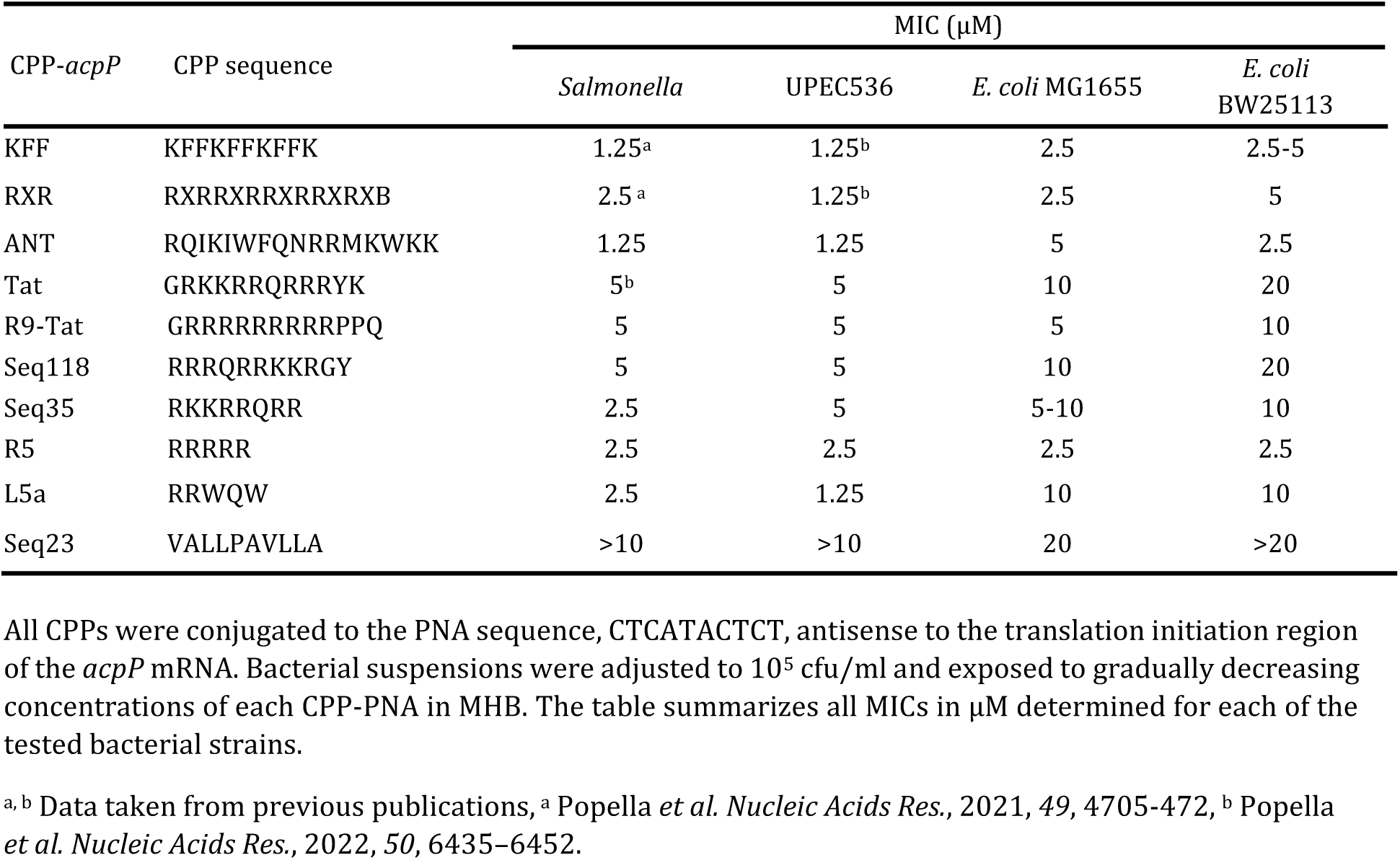
Antibacterial activity of CPP-*acpP* PNAs against *Salmonella* and three *E. coli* strains.

## DISCUSSION

Effective transport across the bacterial envelope remains a bottleneck for the application of antimicrobial ASOs, necessitating the development of new ASO carriers. To facilitate the discovery of better ASO carriers, we developed a “switch-on” reporter system based on antisense-mediated toehold-switch activation (Fig. 1). We show that this system reports ASO uptake in *E. coli* and *Salmonella* and that it can be used to investigate mechanisms of CPP-PNA transport. The switch-on assay presents a simple and semi-quantitative method that lends itself as a building block for future high-throughput screens. Additionally, our work demonstrates that toehold switches can be activated by biostable RNA mimics. This opens their potential application in synthetic biology as tools for orthogonal regulation of multiple genes, each under the control of a different RNA toehold switch, for example for the design of synthetic gene expression networks.

### The switch-on reporter is a versatile tool for high-throughput screens for ASO carriers

ASO uptake into bacteria can be studied using fluorophore labeling of CPP-PNAs (42), mass spectrometry (21, 26), measurement of the antibacterial activity (MIC) of PNAs targeting essential genes (12) or analysis of the transcriptomic responses by using RNA-seq (13, 21). Our assay is a simple alternative that requires no chemical modification of the CPP-PNA (unlike fluorophore labeling) and minimal sample preparation (compared to mass spectrometry and RNA-seq). We validated our reporter assay using two different approaches. First, we showed reduced activation of the toehold switch by KFF-PNA_toe in a bacterial strain lacking the inner membrane transporter SbmA, which is essential for the transport of KFF-PNA conjugates into bacteria (26). Second, we found a good overall correlation of CPP-PNA_toe-mediated increase in sfGFP fluorescence with traditional MIC assays if the identical CPPs were fused to a PNA targeting the essential gene *acpP*. All CPPs that mediated reporter activation showed antibacterial activity, with only one exception (Seq118 in *E. coli* BW25113). This indicates a low probability of predicting false positives, if the reporter assay is used to screen for CPPs for antisense antibiotic delivery. We also observed that L5a was antibacterial although it did not activate the reporter in *Salmonella*, suggesting that other factors beyond ASO delivery can contribute to the antibacterial activity of a CPP-PNA conjugate.

We think that the switch-on reporter assay described here can be developed into a high-throughput screening platform for the discovery of ASO carriers. Bacteria can be treated with a single concentration of each CPP conjugated to PNA_toe (or an optimized version thereof) unlike in MIC determination experiments, which require serial dilutions of the ASO constructs. The fluorescence output can be quantified easily to rank and select CPP based on their efficiency of ASO delivery. The plate reader offers a simple initial read-out but is limited in sensitivity when the fluorescence signal is low. Flow cytometry requires more sample preparation but offers higher sensitivity, which might be important to achieve the dynamic range needed to screen some bacterial strains like the *E. coli* K12 strains used here.

### The toehold switch reporter reveals different dynamics of CPP-mediated PNA delivery

Our screen of ten CPPs revealed different patterns of upregulation in flow cytometry. For example, we observed a rapid but transient increase in fluorescence for some CPPs (e.g., KFF) and a sustained increase for others (e.g., RXR). This variation can likely be explained by the chemistry and stability of these carriers. The KFF peptide is sensitive to proteases and degraded in nutrient-rich media such as MHB (26). By contrast, RXR is composed of a non-canonical 6-aminohexanoic acid and is resistant to proteolytic degradation (43). Therefore, the different dynamics in fluorescence increase could inform about the CPP stability in bacterial culture. Another observed difference among CPPs was that some (KFF, RXR, and ANT) induced strong and rapid upregulation within 4 h, while others, like TAT, were slower, with robust reporter activation only after prolonged incubation. Previously, we have shown that KFF- and RXR-*acpP* eradicate bacteria after 15 minutes while TAT achieves the same effect in 40-60 minutes (13). This indicates that the upregulation of fluorescence from the toehold switch reporter resembles the dynamics of delivery albeit at longer time scales.

### Limitations and optimization of the assay

We established the switch-on reporter assay using a well-studied strain of *Salmonella* as the model and validated its applicability in other enteric bacteria. Our assay shows a high dynamic range in *Salmonella* and UPEC, but the range is more limited in *E. coli* K12 strains. We see two main strategies to enhance the sensitivity and dynamic range of the assay. First, mutations in the stem of the toehold switch will likely increase sensitivity, although these mutations, which partly open the stem loop, bear the risk of higher basal fluorescence. While this might compromise the signal-to-noise ratio, a systematic screen of the TS7 sequence with single or double mutations might nevertheless produce TS7 sequences optimized for application to short ASOs. Such an approach has already been established for the study of 5’ mRNA regions and their regulation by bacterial sRNAs (44). Second, the activator could be improved. Our lead PNA (PNA_toe) achieves a 60-fold activation of the TS7 toehold switch, which is substantially lower than the up to 300-fold activation achieved by the 30mer sRNA in the original work (32). Since strand displacement-assisted activation depends on the length of the competing oligomer (45), longer PNAs might be able to achieve higher activation. However, longer PNAs are usually not internalized efficiently (12, 42). In this study, we have tested two PNAs that target different regions of the stem-loop. PNA_toe was more efficient than PNA_mid, which is consistent with previous reports identifying the toehold as the most crucial site for stem displacement (45). When the activating RNA bound in proximity to the RBS (as does PNA_mid), switch activation was lower (32). This might be caused by interference with ribosome binding, as shown for bacterial sRNAs that repressed translation of target mRNAs by base-pairing just outside the RBS (46, 47). To identify an optimal activator, one could tile the 5’ flank of the TS7 toehold sequence with PNAs of varying lengths, for example by using a recently developed high-throughput synthesis approach in which hundreds of PNA conjugates can be synthesized and then tested *in vivo* (Giorgia Danti, L.P., J.V., Hans Maric., personal communication).

To further improve the assay, independent expression of a different fluorescence protein from the same plasmid could be used to assess potential copy number variations of the reporter plasmid used and bacterial viability. There is a large repertoire of fluorescent proteins that lend themselves as reporters in bacteria, and many of these can be read out in parallel at different wavelengths (48).

While we envision that our work will serve as a template for developing switch-on reporter assays for other pathogens of interest, including Gram-positive bacteria such as *Staphylococcus aureus*, *Enterobacter faecium,* and *Clostridium difficile*, our data show that reporter activation varies between bacterial species, growth media, ASO concentration and design, reporter protein, and the time of analysis post-ASO treatment. Consequently, implementing this assay as a screening platform in other bacteria will require optimization of these factors. For example, some reporter proteins such as GFP do not fold under anaerobic conditions necessary for the growth of some bacterial species, e.g., *C. difficile*; mCherry might be an alternative here.

Recent work in *Fusobacterium nucleatum* has established conditions for the use of different fluorescent proteins under oxygen tension (49), which might help to set up RNA toehold switch assays in a range of other anaerobic bacteria.

### Outlook

Due to the modular nature of CPP-ASOs, precision manipulation of bacterial communities can be achieved either by targeting a species-specific RNA sequence or by using carriers that facilitate selective transport into only one specific species in the community. The assay developed here will enable the high-throughput discovery of novel carriers for the delivery of antisense PNAs, by informing about (i) PNA sequence-independent confounding effects, (ii) uptake kinetics and mechanisms, as well as (iii) species-selectivity of uptake. While we have used our assay to screen CPPs, it may be extended to other classes of carriers, such as nanoparticles (50–53) and siderophores (54, 55). Identifying alternative ASO carriers would allow the use of other ASO backbones such as the negatively charged 2′-4′ carbon bridged-locked nucleic acids (LNA) and 2ʹ-O-methoxyethyl (2ʹ-MOE), which have so far been difficult to deliver with CPPs (21). This could dramatically expand the chemical space for ASOs targeting microbes.

## Materials and methods

### Bacterial strains, PNAs and peptide-conjugated PNAs (CPP-PNAs)

The following strains were used in this study: *Salmonella enterica* serovar Typhimurium strain SL1344 (provided by D. Bumann, MPIIB Berlin, Germany; internal strain number JVS-1574; NCBI GenBank: FQ312003.1), uropathogenic *E. coli* 536 (UPEC 536, internal strain number JVS-12054; NCBI GenBank: CP000247.1), *E. coli* BW25113 (internal strain number JVS-12547; NCBI GenBank: CP009273.1), and *E. coli* MG1655 (internal strain number JVS-5709; NCBI GenBank: U00096.3). All bacterial strains are listed in Supplemental Tables S6.

1 L 5x M9 medium was prepared with 64 g Na_2_HPO_4_.7H_2_O, 15 g KH_2_PO_4_, 2.5 g NaCl, 5 g NH_4_Cl, 2 mM MgSO_4_. 1x M9 medium was diluting 5x M9 in water supplemented with 0.1 mM CaCl_2_, 0.4 % glucose, 0.2 % CAS, 0,004 % Histidine (0,8 %-2,5ml), 25 µl 10 mg/ml Thiamine. Mueller Hinton broth (MHB) purchased from BD Difco™, Thermo Fisher Scientific. Strains were streaked on Luria-Bertani (LB) plates and incubated overnight at 37 °C. Liquid cultures were inoculated in MHB, LB or M9 minimal medium under agitation at 37 °C. Antibiotics were added if needed.

PNAs and peptide-conjugated PNAs (CPP-PNAs) were purchased from Peps4LS GmbH (Heidelberg, Germany). The quality and purity of these constructs was verified by mass spectrometry and HPLC. Unconjugated PNAs and CPP-PNAs (Table S7) were dissolved in water and heated at 55 °C for 5 min before usage. The concentration of the constructs was determined using a NanoDrop spectrophotometer (A_260_ nm). Aliquots of PNAs, CPP-PNAs and peptides were stored at –20 °C and heated at 55 °C for 5 mins before preparing the respective working dilutions. Low-binding pipette tips and Eppendorf tubes (Sarstedt) were used for handling PNAs.

### In vitro transcription

Templates for T7 RNA polymerase-mediated *in vitro* transcription were generated via overlap polymerase chain reaction (PCR) to generate GFP fusion constructs. For this, oligonucleotides were designed to amplify TS7 from an Addgene plasmid (Plasmid #107355), and *gfp* from pXG10 plasmid backbone (34) using a protocol from Popella *et al* (12). The amplicon of TS7 spanned nucleotides –41 to + 30 relative to the translational start codon as described by Green *et al* (32). The PCR for the overlap fusion was performed by using 4 oligonucleotides: 1) Forward primer (JVO-20850) annealing at the 5’ end of TS7 (A T7 promoter sequence was attached to the 5’ end to allow subsequent transcription of the TS7::*gfp* fusion construct *in vitro*) 2) a reverse primer (JVO-20361) amplifying TS7 including a 30 nucleotide *gfp* overlap at the 3’ end (to enable fusion to *gfp*, 3) forward primer annealing at the 5’ end of *gfp* coding sequence (JVO-19762) and 4) reverse primer binding the 3’ end of *gfp* (JVO-19763). All oligonucleotides were purchased from Eurofins Genomics and sequences are listed in Table S4. For PCR amplification, 10 ng of each plasmid was used as a template and all four oligonucleotides were added at a molar ratio of 10:1:1:10 according to Liu et al., 2018 (56). The PCR program was set as follows: 1 min, 98 °C; 40 × cycles of 98 °C for 10 s, 60 °C for 20 s, 72 °C for 40 s; 72 °C for 10 mins, then stored at 4-8 °C until further purification. The PCR product was purified using agarose gel electrophoresis followed by PCR Clean-up (Macherey-Nagel) according to the manufacturer’s instructions and DNA concentration was quantified by using NanoDrop spectrophotometer (A_260_).

After agarose gel-based purification approximately 500 ng of template DNA was subjected to 20 μl *in vitro* transcription reactions following the manufacturer’s instructions (MEGAscript T7 kit, Ambion/Thermo Scientific). The samples were incubated at 37°C for 4 h, after which 2 units (U) of Turbo DNase were added to each reaction for an additional 15 mins. 115 μl water, 15 μl ammonium acetate stop solution, and 3x volume of ethanol were added sequentially to precipitate RNA at –80 °C overnight. The samples were then centrifuged, pellets were washed with 70% ethanol and finally resuspended in 20 μl ultrapure water. RNA concentration was measured with Qubit (Fisher Scientific). RNA gels (6% PAA, 7 M urea) were prepared and stained with StainsAll (Sigma-Aldrich) to verify the expected product size and RNA integrity.

### In vitro translation and Western blotting

To assess PNA-mediated activation of reporter translation *in vitro*, the PURExpress® *In Vitro* Protein Synthesis Kit (New England Biolabs, E6800L) was used according to the manufacturer’s instructions. *In vitro* transcribed RNA was prepared as described above and used in all translation reactions to ensure a precise 100 nM final RNA concentration in each sample. Before starting the translation reaction, TS7::*gfp* RNA was heat-denatured (95 °C for 1 mins), chilled on ice and incubated at 37 °C for 5 mins in the absence or presence of PNA_toe, PNA_ctrl or equal volume of water as negative control. PNA:RNA titrations corresponding to 10:1, 5:1, 2:1, and 1:1 molar ratios were tested. 1 μl of 1 μM RNA was translated in reactions containing 1 μl of 10 µM, 5 µM, 2 µM or 1 μM PNA solutions. Next, PURExpress® *in vitro* translation solution A and B were added and handled as described. Each *in vitro* translation reaction was performed at a final volume of 10 μl, containing 4 μl Solution A, 3 μl Solution B, and 1 pmol of *in vitro* transcribed RNA (adjusted to a final concentration of 100 nM). The reactions were incubated for 2 h at 37 °C after which the tubes were immediately placed on ice. The protein samples were mixed with reducing protein loading buffer (final 1x: 62.6 mM Tris–HCl pH 6.8, 2% SDS, 0.1 mg/ml bromophenol blue, 15.4 mg/ml DTT, 10 % glycerol) and denatured at 95 °C °C for 5 mins. The samples were separated on a SDS-PAA (12% PAA) gel followed by subsequent semi-dry western blot transfer on nitrocellulose membranes to visualize translated protein amounts. The membranes were stained with PonceauS (Sigma-Aldrich) to check for equal loading. Afterwards, membranes were washed, blocked and probed with anti-GFP antibody (1:1000; Sigma-Aldrich) in 5% skim milk (in 1× TBS-T) overnight. After this, the membrane was washed and incubated with an HRP-conjugated secondary antibody (1:10^4^; ThermoScientific) in 1× TBS-T. Protein levels were detected using an ImageQuant LAS 500 (GE Healthcare Life Sciences) following incubation with a self-made developing solution. Images were processed and band intensities were quantified using ImageJ.

### Cloning of translational TS7-reporter plasmids

All plasmids used as PCR or cloning templates or that were generated within this study are listed in Table S5. pNP1 sfGFP toehold switch 7 was a gift from James Collins (Addgene plasmid # 107355; http://n2t.net/addgene:107355; RRID:Addgene_107355). DNA oligonucleotides used for amplification and sequencing were purchased from Eurofins Genomics and are listed in Supplementary Table S4. Oligonucleotide primers were designed to include the NsiI (for forward primer, JVO-21317) and NheI (for reverse primer, JVO-21318) to amplify TS7 (–41 to + 30 relative to the translational start codon) from the Addgene plasmid (Plasmid #107355). The TS7 insert was generated using the PCR program as follows: 1 min, 98 °C; 40 × cycles of 98 °C for 10 s, 58 °C for 20 s, 72 °C for 40 s; 72 °C for 10 mins, then stored at 4-8 °C until further purification. The PCR product of the correct size (118 bp) was purified with 1% agarose gel electrophoresis followed by PCR Clean-up (Macherey-Nagel) according to the manufacturer’s instructions. The pXG10-SF backbone (34) as well as the TS7 insert were cut with NsiI/NheI for subsequent cloning. For the *TS7::mCherry* construct, *mCherry* was PCR amplified (JVO-19329/19330) and cloned into TS7 containing pXG10-SF (pPS002 backbone), by restriction cloning at NheI/XbaI sites. For the *TS7::mNeonGreen* construct, fusion PCR was used to generate the inserts (JVO-21317/21512/JVO-21511/20632) including NheI and XbaI restriction sites. The vector pXG10-SF (34) was cut at NsiI/XbaI and TS7::*mNeonGreen* was inserted using restriction cloning. Single nucleotide mutagenesis for the construction of mutated toehold switches, TS7-Mut_9, TS7-Mut_13, and TS7-Mut_18 were performed on pPS002 using primer pairs specified in Table S5. Briefly, mutations were generated using the PCR program as follows: 30 s, 98 °C; 40 × cycles of 98 °C for 10 s, 52 °C for 30 s, 72 °C for 2 mins; 72 °C for 10 min, then stored at 4-8 °C until further purification. The plasmid was purified with 1% agarose gel electrophoresis followed by PCR Clean-up (Macherey-Nagel) according to the manufacturer’s instructions. Plasmids were sequenced and validated using primers listed in Table S4. *E. coli* TOP10 (Invitrogen) was used for all cloning procedures.

### CPP-PNA treatment of bacteria for sfGFP measurement

Overnight cultures of respective bacterial strains (listed in Table S7) were diluted 1:100 in fresh MHB and grown to OD_600_ 0.3. Overnight and secondary cultures were prepared with or without chloramphenicol (20 µg/ml) depending on the antibiotic resistance profile of the strain. 95 µl of the 0.3 OD bacterial suspension were distributed in the 96-well plate, mixed with 5 µl water or 20× CPP-PNA working solutions (150 µM, 100 µM or 50 µM stocks in water). An equivalent volume of water was added to the bacteria to serve as untreated control. For testing the activity in other growth media (M9 minimal medium and LB), the bacteria were cultured in the respective media and treated as described above.

The PNA-treated samples were incubated in the plate reader for 4-24 hours at 37°C, with continuous double-orbital shaking (237 cpm). The fluorescence (λ_ex_ 480 nm/ λ_em_ 510 nm) and OD_600_ were recorded in a Synergy H1 plate reader (Biotek) by measuring every 20 mins. The bacteria were sampled at various time points post-treatment and further analyzed by flow cytometry.

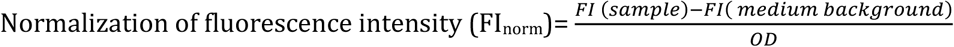

Fold upregulation in FI from the plate reader was calculated as:

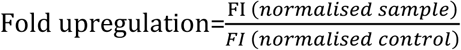

### Flow cytometric measurement and data analysis

Bacterial cells were treated as described above, fixed with 4% PFA in phosphate-buffered saline (PBS) after the indicated time points and stained with 1 μg/ml DAPI for flow cytometric measurement of DAPI-positive bacterial cells. Cell suspensions were diluted by a factor of ∼100-1,000 in PBS and sampled from 96-well plates to achieve ∼2,000 counts/sec for flow cytometric measurements. Flow cytometry was performed using the Agilent Novocyte flow cytometer with a high-throughput sampler and 100,000 events were recorded per sample. The DAPI fluorescence was detected using the Pacific Blue channel; sfGFP and mNeon were detected in the FITC channel, and mCherry fluorescence in the PE-Texas Red channel. The FSC-H threshold was set to 2,000. The wild-type strains without plasmids served as a control to detect the background fluorescence, termed bacterial background. A contour plot with FSC and SSC, each with bins defined on a logarithmic scale was used to select singlets. The gating strategy is described in Fig. S5. The median fluorescence intensity was used for calculating the fold upregulation of fluorescence. Flow cytometry datasets were analyzed using the Novocyte software (Agilent).

Fold upregulation in median fluorescence intensity (FI) from flow cytometry was calculated as:

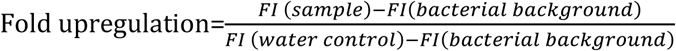

The fold upregulation was calculated from at least 2 independent experiments.

### Minimal inhibitory concentration (MIC) assay

The broth microdilution method was applied for the determination of MIC values according to a previously modified protocol based on the Clinical and Laboratory Standards Institute guidelines (35, 57). Overnight bacterial cultures were diluted 100-fold in fresh MHB and grown to an OD_600_ 0.5. The obtained culture was diluted in fresh MHB to adjust a final cell concentration of ∼10^5^ cfu/mL. 95 µL of the diluted bacterial solution was dispensed into a 96-well plate (Thermo Fisher Scientific). 5 µL of 20x CPP-PNA working solutions (ranging from 400 to 6.25 µM) or an equivalent volume of water as a negative control was added (Table S7). Growth (OD_600_) was monitored in a Synergy H1 plate reader (Biotek) by measuring every 20 mins with continuous double-orbital shaking (237 cpm) at 37 °C for 24 h. The MIC was determined as the lowest concentration of CPP-PNA, which inhibited visible growth in the wells (OD < 0.05).

## Author contributions

P.S. designed the research, performed most of the experiments with the support of L.P. and S.P.J., analyzed the data, and prepared the original draft and wrote the manuscript; L.P. and S.P.J., validated results, reviewed and edited the manuscript; J.V. acquired funding, conceptualized the project, and wrote the manuscript.

## Acknowledgements

We are grateful to Dr. Anke Sparmann for editing the manuscript and to Barbara Plaschke and Phung Thao Do for their technical assistance. We thank Prof. Chase Beisel for fruitful scientific discussions. Research was funded by the Bavarian Bayresq.net (L.P., J.V., P.S.), the BMBF in the framework of the Cluster4Future program (Cluster for Nucleic Acid Therapeutics Munich, CNATM (Project ID: 03ZU1201CA, L.P., J.V.), and a Leibniz Award by the German Research Council (to J.V.; DFG Vo875/18).

## Supplemental information

**Figure S1.**
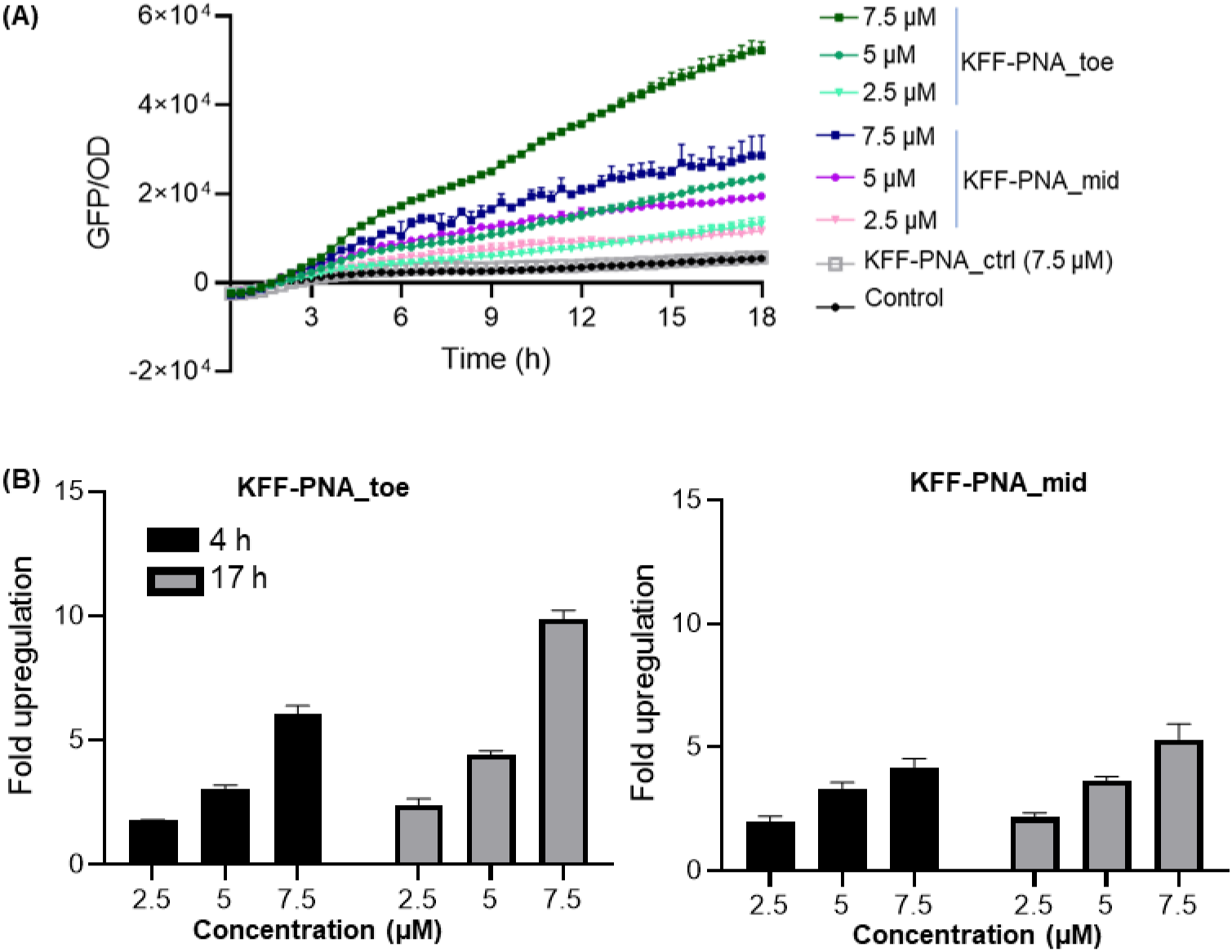
Assessment of fluorescence induction in *Salmonella* (transformed with a plasmid expressing *TS7::sfGFP*) post-treatment with KFF-PNAs targeting different regions of the toehold switch. (A) Normalized fluorescence intensity over time and (B) fold-upregulation of fluorescence of bacteria treated with KFF-PNA_toe and PNA_mid, respectively, measured in a microplate reader. Bacteria were cultured to an OD_600_ of 0.3 and treated with 2.5, 5, and 7.5 μM concentrations of each KFF-PNA. A non-targeting KFF-PNA (PNA_ctrl) was tested to assess non-specific upregulation from KFF-PNA at the highest concentration (7.5 μM). ‘Control’ denotes treatment with an equal volume of water. The fluorescence intensity obtained from the microplate reader was normalized relative to the OD_600_ to account for the decrease in fluorescence due to growth retardation. Bars indicate fold upregulation of PNA-treated samples relative to the water control and error bars indicate the standard deviation calculated from two independent experiments.

**Figure S2.**
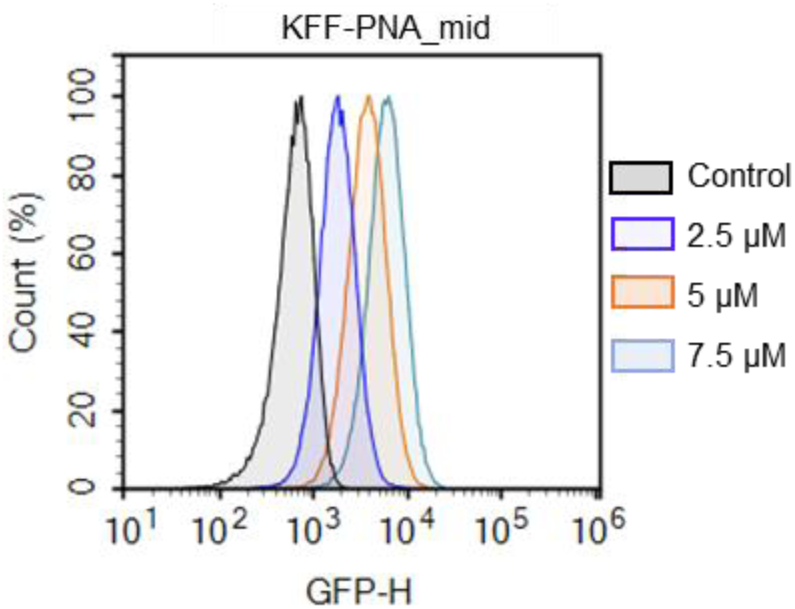
Flow cytometric analysis of activation of *TS7::sfGFP* with KFF-PNA_mid. The flow cytometry histogram represents mean fluorescence intensity of sfGFP 4 h post-treatment of *Salmonella*. *Salmonella* were cultured to an OD_600_ of 0.3 and treated with 2.5, 5, and 7.5 μM concentrations of KFF-PNA_mid. The water-treated sample served as the control condition (Control).

**Figure S3.**
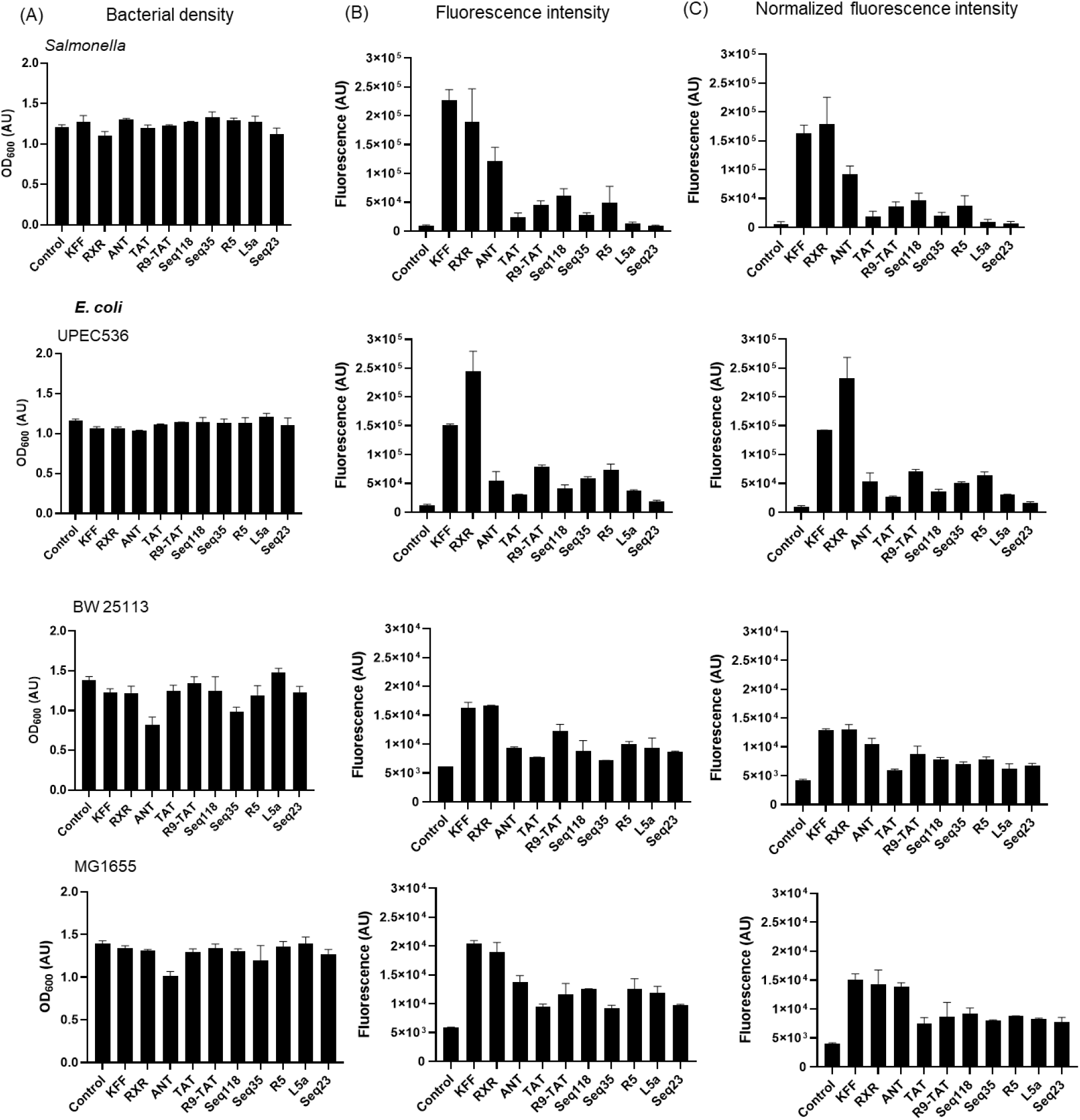
Fluorescence intensity is normalized to OD to enable comparison of PNA delivery efficacy. (A) OD_600_, (B) raw fluorescence intensity and (C) fluorescence intensity normalized to OD_600_ of bacteria expressing *TS7-Mut_9::sfgfp* constructs, 17 h post-treatment with 5 µM concentrations of CPP-PNA_toe measured in a microplate reader. Bacteria were cultured to an OD_600_ of 0.3 and treated with CPP-PNA. The water-treated sample served as control. The fluorescence intensity obtained from the microplate reader was normalized relative to the OD_600_ to account for the decrease in fluorescence due to growth retardation. Bars indicate the mean fluorescence intensity of PNA-treated and control samples. Error bars indicate the standard deviation of mean fold upregulation calculated from two independent experiments.

**Figure S4.**
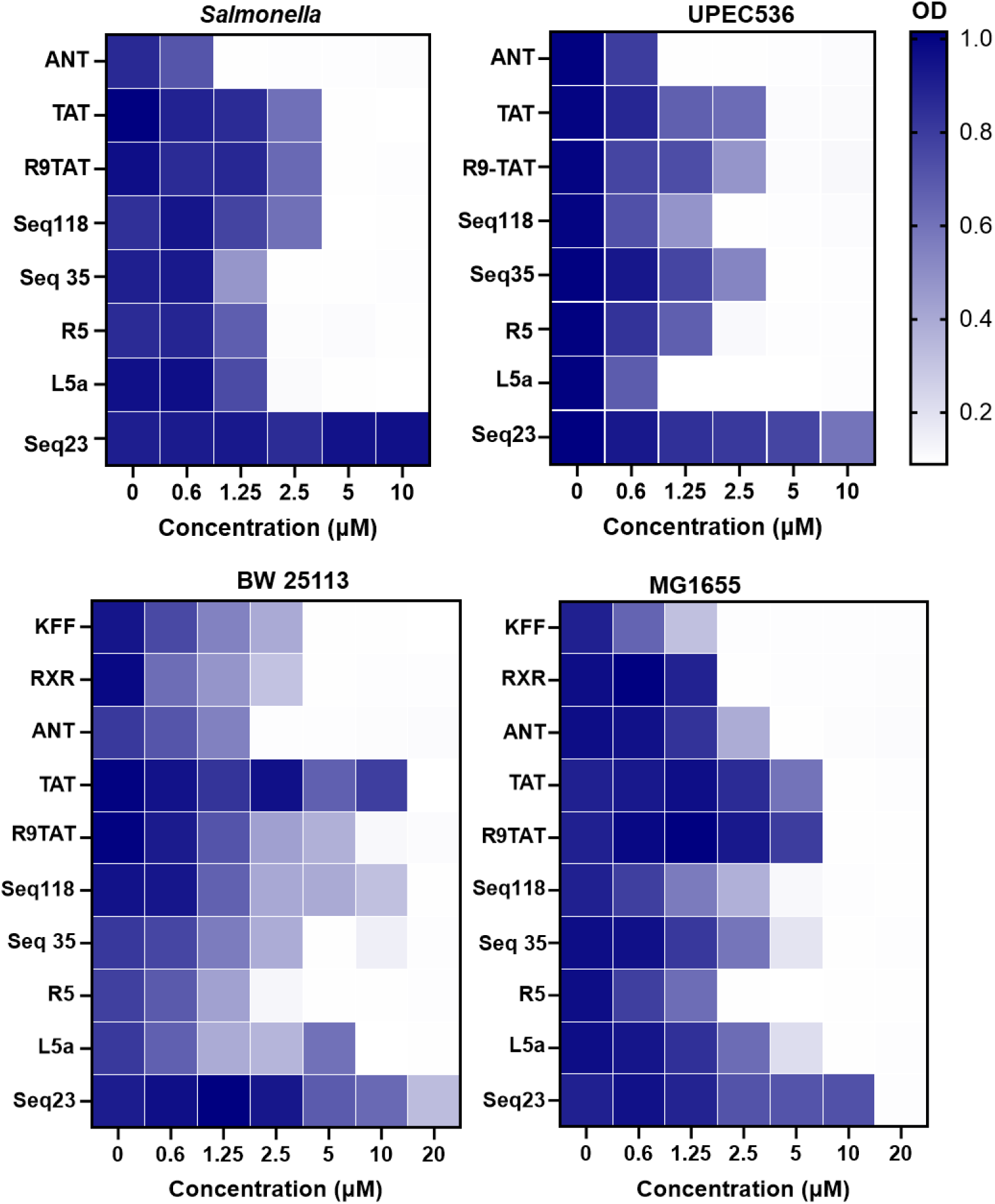
MIC determination validates the reliability of the switch-on reporter assay. The antibacterial activity of the 10 CPPs when conjugated to an anti-*acpP* PNA was tested against various *wt* strains of *Salmonella* and *E. coli.* The heatmaps indicate the OD_600_ of bacteria 20 h post-treatment with varying concentrations of CPP-PNAs. Row titles indicate CPP; all CPPs were conjugated to the PNA sequence CTCATACTCT, antisense to the translation initiation region of the *acpP* mRNA. Column titles indicate the concentration of CPP-*acpP*.

**Figure S5.**
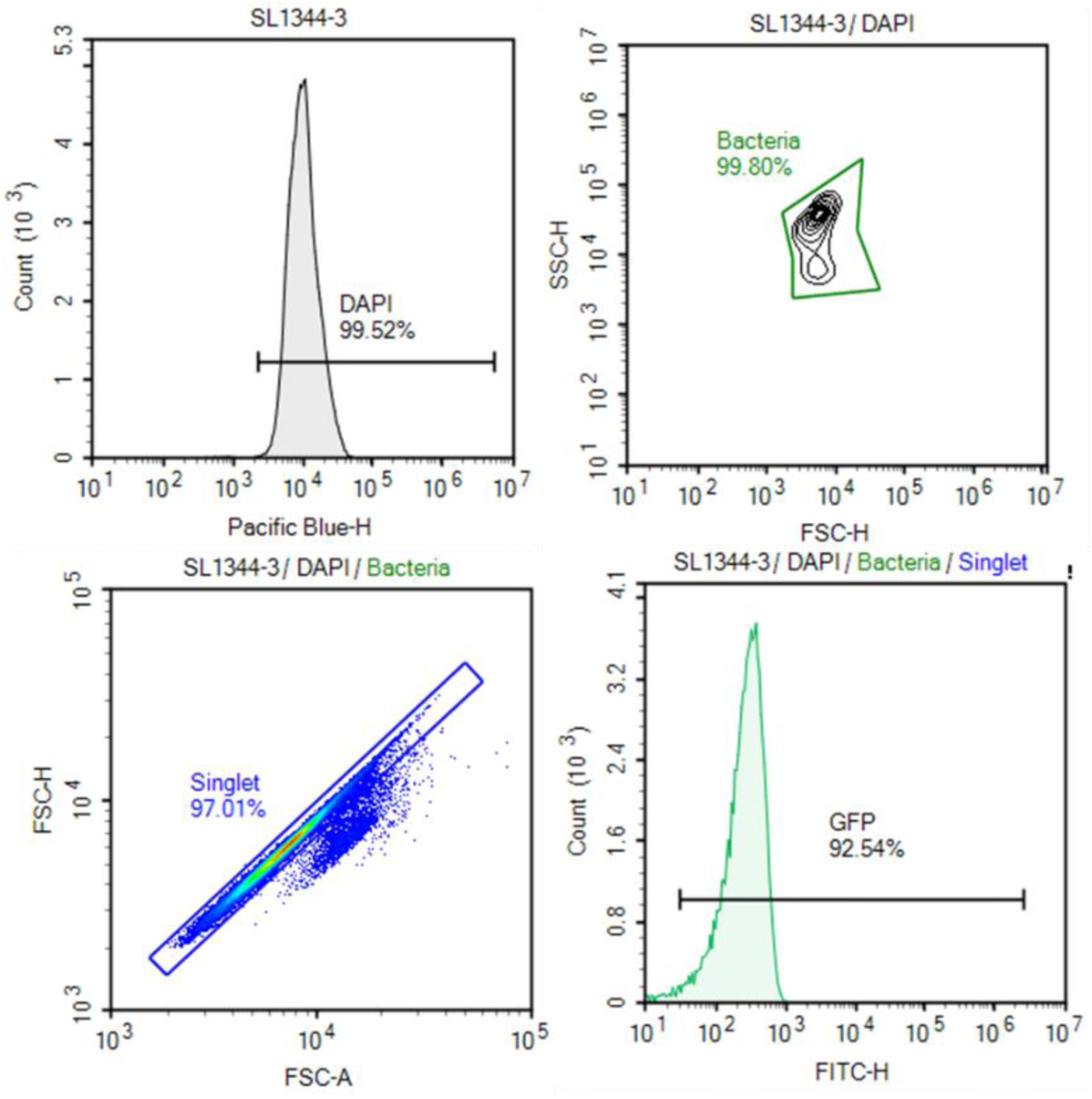
Gating strategy for flow cytometry. *Salmonella* were fixed and stained with DAPI to include only intact cells and 100,000 events were recorded in the Pacific Blue channel. The fluorescence of DAPI was recorded using the Pacific Blue channel; sfGFP was detected in the FITC channel. The FSC-H threshold was set to 2000.

---

**Table S1.**
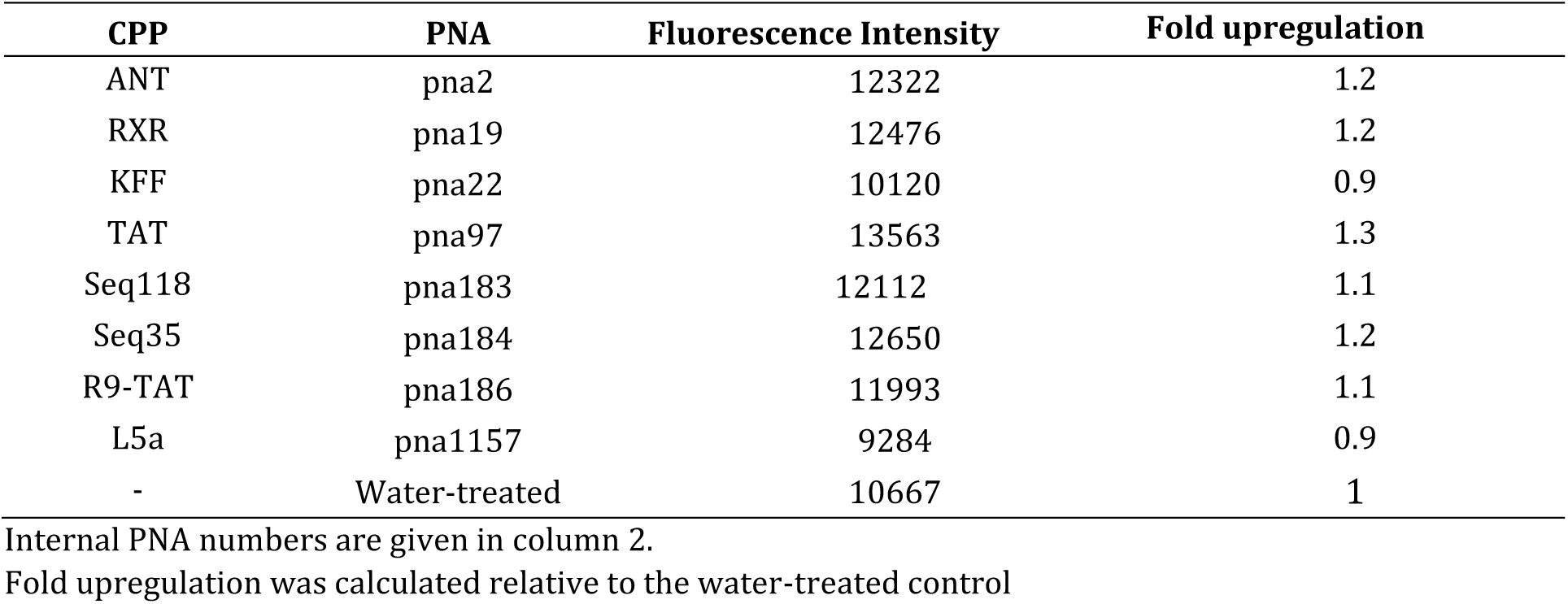
Fluorescence intensity of *Salmonella* carrying plasmids encoding *TS7-Mut_9::sfgfp*, 17 h post-treatment with a non-targeting PNA sequence (PNA_ctrl) conjugated with various CPPs measured using a microplate reader.

| CPP | PNA | Fluorescence Intensity | Fold upregulation |
| --- | --- | --- | --- |
| ANT | pna2 | 12322 | 1.2 |
| RXR | pna19 | 12476 | 1.2 |
| KFF | pna22 | 10120 | 0.9 |
| TAT | pna97 | 13563 | 1.3 |
| Seq118 | pna183 | 12112 | 1.1 |
| Seq35 | pna184 | 12650 | 1.2 |
| R9-TAT | pna186 | 11993 | 1.1 |
| L5a | pna1157 | 9284 | 0.9 |
| - | Water-treated | 10667 | 1 |

Internal PNA numbers are given in column 2.
Fold upregulation was calculated relative to the water-treated control

**Table S2.** Median fluorescence intensity of *Salmonella* and *E. coli* expressing *TS7::sfGFP*, at various time points post-treatment with non-specific PNA, PNA_ctrl sequence conjugated with various CPPs measured using flow cytometry.

| Sample | Median Fluorescence (GFP-H) |  |  |  |
| --- | --- | --- | --- | --- |
|  | *Salmonella* | *E. coli* MG1655 | *E. coli* BW25113 | UPEC536 |
| **4 h** |  |  |  |  |
| ANT | 523 | 554 | 564 | 650 |
| RXR | 488 | 537 | 585 | 466 |
| KFF | 491 | 549 | 577 | 634 |
| TAT | 537 | 549 | 582 | 643 |
| Seq118 | 526 | 553 | 585 | 635 |
| Seq35 | 509 | 549 | 596 | 639 |
| R9-TAT | 476 | 570 | 610 | 646 |
| L5a | 470 | 553 | 602 | 643 |
| Water control* | 479 | 554 | 601 | 634 |
| **6 h** |  |  |  |  |
| ANT | 564 | 481 | 519 | 662 |
| RXR | 445 | 471 | 490 | 420 |
| KFF | 507 | 482 | 498 | 642 |
| TAT | 595 | 476 | 499 | 658 |
| Seq118 | 537 | 477 | 501 | 642 |
| Seq35 | 527 | 479 | 510 | 653 |
| R9-TAT | 531 | 423 | 515 | 646 |
| L5a | 542 | 487 | 499 | 650 |
| Water control* | 500 | 488 | 503 | 644 |
| **17 h** |  |  |  |  |
| ANT | 748 | 548 | 488 | 926 |
| RXR | 472 | 540 | 494 | 596 |
| KFF | 669 | 542 | 458 | 835 |
| TAT | 716 | 556 | 444 | 896 |
| Seq118 | 594 | 545 | 440 | 1099 |
| Seq35 | 641 | 567 | 476 | 993 |
| R9-TAT | 530 | 560 | 455 | 911 |
| L5a | 661 | 554 | 443 | 953 |
| Water control* | 656 | 548 | 447 | 963 |

\*Water control denotes the bacterial sample treated with only an equivalent volume of water, serving as the baseline.

**Table S4.**
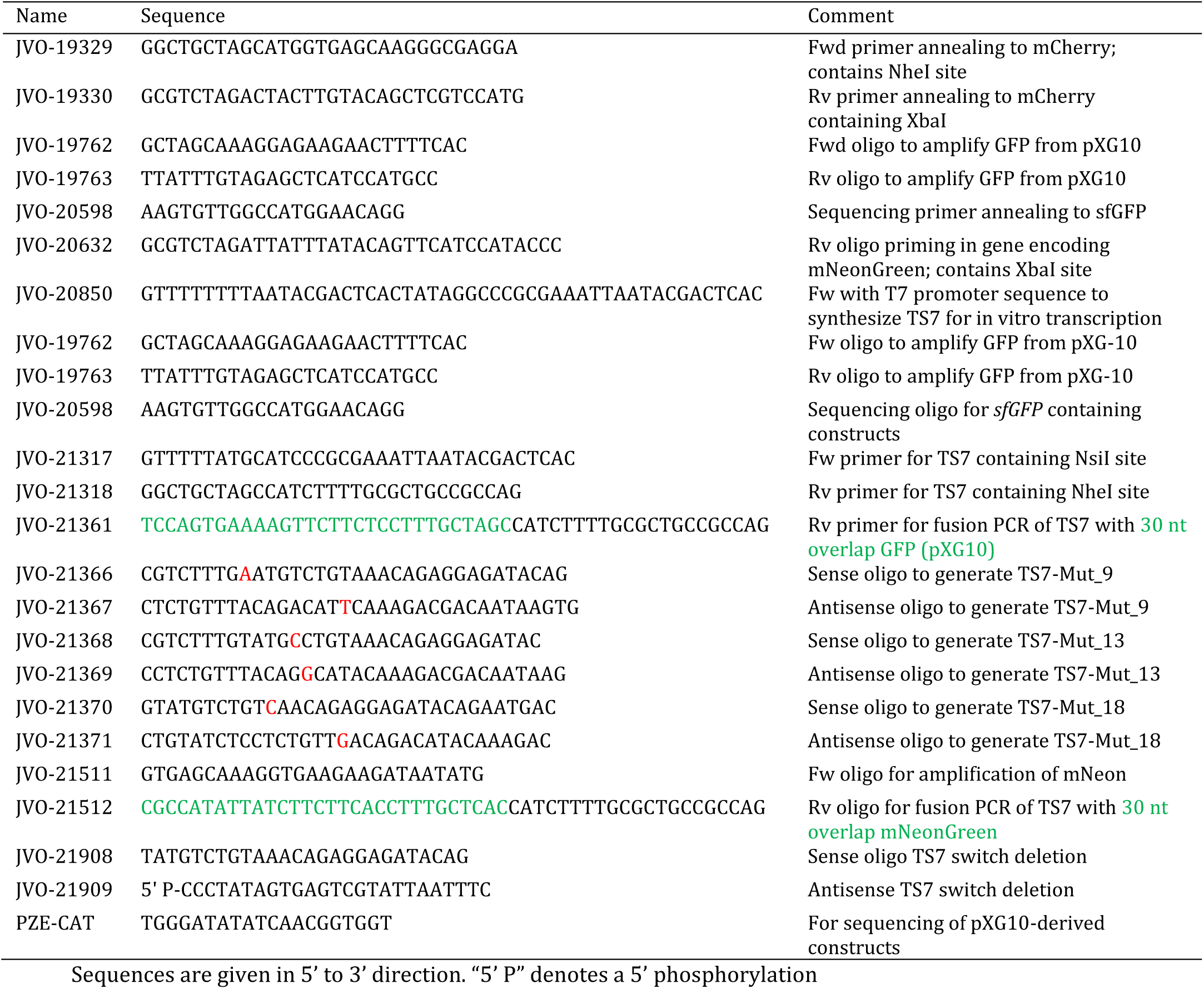
List of oligonucleotides used in this study.

**Table S5.** List of plasmids used in this study.

| Plasmid Number | Insert name | Backbone/Marker | Oligos for insert amplification | Cloning sites | Description |
| --- | --- | --- | --- | --- | --- |
| pPS002 | TS7 | pXG10-sf/Cm | JVO21317/<br>JVO21318 | NsiI/NheI | TS7 insert was cloned from pNP1-TS7 obtained from Addgene |
| pPS007 | TS7-sfGFP | pNP1/Amp |  |  | Addgene Plasmid #107355 |
| pPS013 | TS7-Mut_18 | pXG10-SF/Cm | JVO21370/<br>JVO21371 |  | Mutations induced into pXG10-TS7 (pPS2) at pos 18 of stem |
| pPS014 | TS7 | pXG10/Cm | JVO-21317/<br>JVO-20632/<br>JVO-21511/<br>JVO-21512/ | NheI/XbaI | Template TS7: pPS002<br>Fused to mNeon (from pKAS32mNeon) followed by cloning into in pXG10sf |
| pPS016 | TS7-Mut_9 | pXG10-SF/Cm | JVO21366/<br>JVO21367 |  | Single site mutation induced into TS7 |
| pPS017 | TS7-Mut_13 | pXG10-SF/Cm | JVO21368/<br>JVO21369 |  | Single site mutation induced into TS7 |
| pPS020 | TS7-mCherry | pXG10-mCherry/Cm | JVO19329/<br>JVO 19330 | NheI/ XbaI | Template pSDM007<br>Backbone pPS002 |
| pPS035 | TS7-Mut_9 | pXG10-mNeon/Cm | JVO21366/<br>JVO21367 | Cm | Template pPS014 |
| pPS036 | TS7-Mut_9 | pXG10-mCherry/Cm | JVO21366/<br>JVO21367 | Cm | Template pPS020 |

Template indicates template used for insert synthesis. Backbone indicates vector.

**Table S6.**
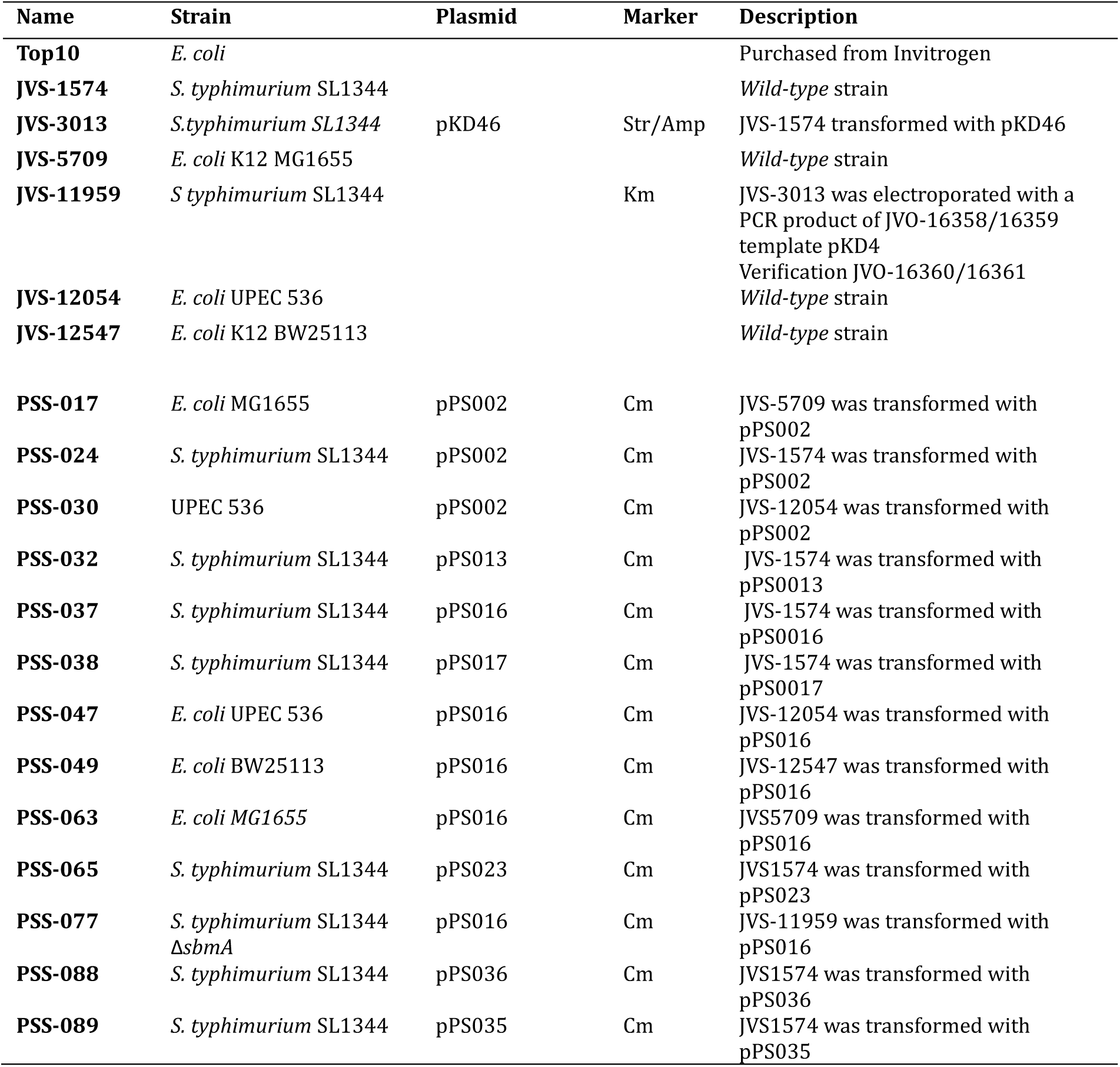
List of strains used in this study.

**Table S7.**
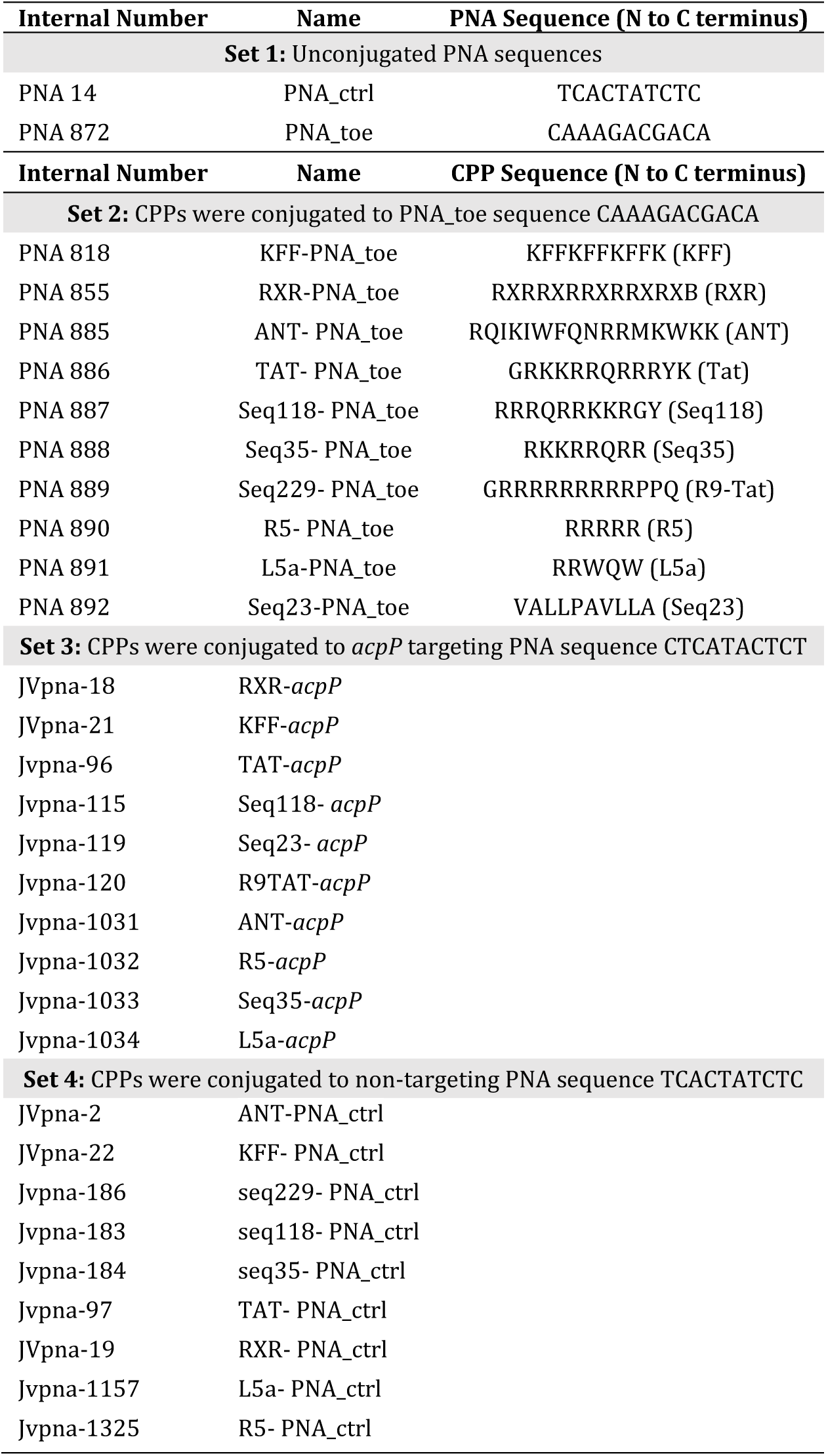
List of CPP-PNAs used in this study.

## References

1. Antimicrobial Resistance Collaborators. 2022. Global burden of bacterial antimicrobial resistance in 2019: a systematic analysis. Lancet 399:629–655.

2. Pifer R, Greenberg DE. 2020. Antisense antibacterial compounds. Transl Res 223:89–106.

3. Vogel J. 2020. An RNA biology perspective on species-specific programmable RNA antibiotics. Mol Microbiol 113:550–559.

4. El-Fateh M, Chatterjee A, Zhao X. 2024. A systematic review of peptide nucleic acids (PNAs) with antibacterial activities: Efficacy, potential and challenges. Int J Antimicrob Agents 63:107083.

5. Dryselius R, Aswasti SK, Rajarao GK, Nielsen PE, Good L. 2003. The translation start codon region is sensitive to antisense PNA inhibition in *Escherichia coli*. Oligonucleotides 13:427–33.

6. Good L, Awasthi SK, Dryselius R, Larsson O, Nielsen PE. 2001. Bactericidal antisense effects of peptide-PNA conjugates. Nat Biotechnol 19:360–4.

7. Otoupal PB, Eller KA, Erickson KE, Campos J, Aunins TR, Chatterjee A. 2021. Potentiating antibiotic efficacy via perturbation of non-essential gene expression. Commun Biol 4:1267.

8. Mu Y, Shen Z, Jeon B, Dai L, Zhang Q. 2013. Synergistic effects of anti-CmeA and anti-CmeB peptide nucleic acids on sensitizing Campylobacter jejuni to antibiotics. Antimicrob Agents Chemother 57:4575–7.

9. Castillo JI, Rownicki M, Wojciechowska M, Trylska J. 2018. Antimicrobial synergy between mRNA targeted peptide nucleic acid and antibiotics in E. coli. Bioorg Med Chem Lett 28:3094–3098.

10. Goh S, Loeffler A, Lloyd DH, Nair SP, Good L. 2015. Oxacillin sensitization of methicillin-resistant Staphylococcus aureus and methicillin-resistant *Staphylococcus pseudintermedius* by antisense peptide nucleic acids in vitro. BMC Microbiol 15:262.

11. Ayhan DH, Tamer YT, Akbar M, Bailey SM, Wong M, Daly SM, Greenberg DE, Toprak E. 2016. Sequence-Specific Targeting of Bacterial Resistance Genes Increases Antibiotic Efficacy. PLoS Biol 14:e1002552.

12. Popella L, Jung J, Do PT, Hayward RJ, Barquist L, Vogel J. 2022. Comprehensive analysis of PNA-based antisense antibiotics targeting various essential genes in uropathogenic Escherichia coli. Nucleic Acids Res 50:6435–6452.

13. Popella L, Jung J, Popova K, Ethurica-Mitic S, Barquist L, Vogel J. 2021. Global RNA profiles show target selectivity and physiological effects of peptide-delivered antisense antibiotics. Nucleic Acids Res 49:4705–4724.

14. Hizume T, Sato Y, Iwaki H, Honda K, Okano K. 2023. Subtractive modification of bacterial consortium using antisense peptide nucleic acids. Front Microbiol 14:1321428.

15. Moustafa DA, Wu AW, Zamora D, Daly SM, Sturge CR, Pybus C, Geller BL, Goldberg JB, Greenberg DE. 2021. Peptide-Conjugated Phosphorodiamidate Morpholino Oligomers Retain Activity against Multidrug-Resistant Pseudomonas aeruginosa In Vitro and In Vivo. mBio 12.

16. Kurupati P, Tan KS, Kumarasinghe G, Poh CL. 2007. Inhibition of gene expression and growth by antisense peptide nucleic acids in a multiresistant beta-lactamase-producing *Klebsiella pneumoniae* strain. Antimicrob Agents Chemother 51:805–11.

17. Jung J, Popella L, Do PT, Pfau P, Vogel J, Barquist L. 2023. Design and off-target prediction for antisense oligomers targeting bacterial mRNAs with the MASON web server. RNA 29:570–583.

18. Daly SM, Sturge CR, Marshall-Batty KR, Felder-Scott CF, Jain R, Geller BL, Greenberg DE. 2018. Antisense Inhibitors Retain Activity in Pulmonary Models of Burkholderia Infection. ACS Infect Dis 4:806–814.

19. Geller BL, Marshall-Batty K, Schnell FJ, McKnight MM, Iversen PL, Greenberg DE. 2013. Gene-silencing antisense oligomers inhibit acinetobacter growth in vitro and in vivo. J Infect Dis 208:1553–60.

20. Barkowsky G, Abt C, Pohner I, Bieda A, Hammerschmidt S, Jacob A, Kreikemeyer B, Patenge N. 2022. Antimicrobial Activity of Peptide-Coupled Antisense Peptide Nucleic Acids in Streptococcus pneumoniae. Microbiol Spectr 10:e0049722.

21. Ghosh C, Popella L, Dhamodharan V, Jung J, Dietzsch J, Barquist L, Hobartner C, Vogel J. 2024. A comparative analysis of peptide-delivered antisense antibiotics using diverse nucleotide mimics. RNA 30:624–643.

22. Hadjicharalambous A, Bournakas N, Newman H, Skynner MJ, Beswick P. 2022. Antimicrobial and Cell-Penetrating Peptides: Understanding Penetration for the Design of Novel Conjugate Antibiotics. Antibiotics (Basel) 11.

23. Kazusato Oikawa MMI, Yoko Horii, Takeshi Yoshizumi, and Keiji Numata*. 2018. Screening of a Cell-Penetrating Peptide Library in Escherichia coli: Relationship between Cell Penetration Efficiency and Cytotoxicity. ACS Omega 12:16489–16499.

24. Lee HM, Ren J, Tran KM, Jeon BM, Park WU, Kim H, Lee KE, Oh Y, Choi M, Kim DS, Na D. 2021. Identification of efficient prokaryotic cell-penetrating peptides with applications in bacterial biotechnology. Commun Biol 4:205.

25. Jha D, Mishra R, Gottschalk S, Wiesmuller KH, Ugurbil K, Maier ME, Engelmann J. 2011. CyLoP-1: a novel cysteine-rich cell-penetrating peptide for cytosolic delivery of cargoes. Bioconjug Chem 22:319–28.

26. Yavari N, Goltermann L, Nielsen PE. 2021. Uptake, Stability, and Activity of Antisense Anti-acpP PNA-Peptide Conjugates in Escherichia coli and the Role of SbmA. ACS Chem Biol 16:471–479.

27. Meng J, Da F, Ma X, Wang N, Wang Y, Zhang H, Li M, Zhou Y, Xue X, Hou Z, Jia M, Luo X. 2015. Antisense growth inhibition of methicillin-resistant Staphylococcus aureus by locked nucleic acid conjugated with cell-penetrating peptide as a novel FtsZ inhibitor. Antimicrob Agents Chemother 59:914–22.

28. Shao Y, Wu Y, Chan CY, McDonough K, Ding Y. 2006. Rational design and rapid screening of antisense oligonucleotides for prokaryotic gene modulation. Nucleic Acids Res 34:5660–9.

29. Tilley LD, Hine OS, Kellogg JA, Hassinger JN, Weller DD, Iversen PL, Geller BL. 2006. Gene-specific effects of antisense phosphorodiamidate morpholino oligomer-peptide conjugates on *Escherichia coli* and *Salmonella enterica* serovar typhimurium in pure culture and in tissue culture. Antimicrob Agents Chemother 50:2789–96.

30. Wierzba AJ, Wojciechowska M, Trylska J, Gryko D. 2021. Vitamin B(12) - Peptide Nucleic Acid Conjugates. Methods Mol Biol 2355:65–82.

31. Papenfort K, Vanderpool CK. 2015. Target activation by regulatory RNAs in bacteria. FEMS Microbiol Rev 39:362–78.

32. Green AA, Silver PA, Collins JJ, Yin P. 2014. Toehold switches: de-novo-designed regulators of gene expression. Cell 159:925–39.

33. Jasinski M, Miszkiewicz J, Feig M, Trylska J. 2019. Thermal Stability of Peptide Nucleic Acid Complexes. J Phys Chem B 123:8168–8177.

34. Corcoran CP, Podkaminski D, Papenfort K, Urban JH, Hinton JC, Vogel J. 2012. Superfolder GFP reporters validate diverse new mRNA targets of the classic porin regulator, MicF RNA. Mol Microbiol 84:428–45.

35. Performance Standards for Antimicrobial Susceptibility Testing M100, CLSI Guidelines. https://clsi.org/standards/products/microbiology/.

36. Robinson JP, Ostafe R, Iyengar SN, Rajwa B, Fischer R. 2023. Flow Cytometry: The Next Revolution. Cells 12.

37. Ehrenberg M, Bremer H, Dennis PP. 2013. Medium-dependent control of the bacterial growth rate. Biochimie 95:643–58.

38. Bock LJ, Hind CK, Sutton JM, Wand ME. 2018. Growth media and assay plate material can impact on the effectiveness of cationic biocides and antibiotics against different bacterial species. Lett Appl Microbiol 66:368–377.

39. Pedelacq JD, Cabantous S, Tran T, Terwilliger TC, Waldo GS. 2006. Engineering and characterization of a superfolder green fluorescent protein. Nat Biotechnol 24:79–88.

40. Balleza E, Kim JM, Cluzel P. 2018. Systematic characterization of maturation time of fluorescent proteins in living cells. Nat Methods 15:47–51.

41. Liu BR, Huang YW, Aronstam RS, Lee HJ. 2016. Identification of a Short Cell-Penetrating Peptide from Bovine Lactoferricin for Intracellular Delivery of DNA in Human A549 Cells. PLoS One 11:e0150439.

42. Goltermann L, Yavari N, Zhang M, Ghosal A, Nielsen PE. 2019. PNA Length Restriction of Antibacterial Activity of Peptide-PNA Conjugates in Escherichia coli Through Effects of the Inner Membrane. Front Microbiol 10:1032.

43. Mellbye BL, Puckett SE, Tilley LD, Iversen PL, Geller BL. 2009. Variations in amino acid composition of antisense peptide-phosphorodiamidate morpholino oligomer affect potency against *Escherichia coli* in vitro and in vivo. Antimicrob Agents Chemother 53:525–30.

44. Holmqvist E, Reimegard J, Wagner EG. 2013. Massive functional mapping of a 5’-UTR by saturation mutagenesis, phenotypic sorting and deep sequencing. Nucleic Acids Res 41:e122.

45. Simmel FC, Yurke B, Singh HR. 2019. Principles and Applications of Nucleic Acid Strand Displacement Reactions. Chem Rev 119:6326–6369.

46. Bouvier M, Sharma CM, Mika F, Nierhaus KH, Vogel J. 2008. Small RNA binding to 5’ mRNA coding region inhibits translational initiation. Mol Cell 32:827–37.

47. Sharma CM, Darfeuille F, Plantinga TH, Vogel J. 2007. A small RNA regulates multiple ABC transporter mRNAs by targeting C/A-rich elements inside and upstream of ribosome-binding sites. Genes Dev 21:2804–17.

48. Delgadillo-Guevara M, Halte M, Erhardt M, Popp PF. 2024. Fluorescent tools for the standardized work in Gram-negative bacteria. J Biol Eng 18:25.

49. Ponath F, Zhu Y, Cosi V, Vogel J. 2022. Expanding the genetic toolkit helps dissect a global stress response in the early-branching species *Fusobacterium nucleatum*. Proc Natl Acad Sci U S A 119:e2201460119.

50. Moreira L, Guimaraes NM, Pereira S, Santos RS, Loureiro JA, Pereira MC, Azevedo NF. 2022. Liposome Delivery of Nucleic Acids in Bacteria: Toward In Vivo Labeling of Human Microbiota. ACS Infect Dis 8:1218–1230.

51. Pereira S, Santos RS, Moreira L, Guimaraes N, Gomes M, Zhang H, Remaut K, Braeckmans K, De Smedt S, Azevedo NF. 2021. Lipoplexes to Deliver Oligonucleotides in Gram-Positive and Gram-Negative Bacteria: Towards Treatment of Blood Infections. Pharmaceutics 13:989.

52. Li A, Zhang Y, Wan L, Peng R, Zhang X, Guo Q, Xu S, Qiao D, Zheng P, Li N, Zhu W, Pan Q. 2024. Coordination-Driven Self-Assembly of Metal Ion-Antisense Oligonucleotide Nanohybrids for Chronic Bacterial Infection Therapy. ACS Appl Mater Interfaces 16:28041–28055.

53. Liu M, Chu B, Sun R, Ding J, Ye H, Yang Y, Wu Y, Shi H, Song B, He Y, Wang H, Hong J. 2023. Antisense Oligonucleotides Selectively Enter Human-Derived Antibiotic-Resistant Bacteria through Bacterial-Specific ATP-Binding Cassette Sugar Transporter. Adv Mater 35:e2300477.

54. Pals MJ, Wijnberg L, Yildiz C, Velema WA. 2024. Catechol-Siderophore Mimics Convey Nucleic Acid Therapeutics into Bacteria. Angew Chem Int Ed Engl 63:e202402405.

55. Tsylents U, Burmistrz M, Wojciechowska M, Stepien J, Maj P, Trylska J. 2024. Iron uptake pathway of *Escherichia coli* as an entry route for peptide nucleic acids conjugated with a siderophore mimic. Front Microbiol 15:1331021.

56. Liu Y, Chen J, Thygesen A. 2018. Efficient One-Step Fusion PCR Based on Dual-Asymmetric Primers and Two-Step Annealing. Mol Biotechnol 60:92–99.

57. Goltermann L, Nielsen PE. 2020. PNA Antisense Targeting in Bacteria: Determination of Antibacterial Activity (MIC) of PNA-Peptide Conjugates. Methods Mol Biol 2105:231–239.

